# Chandipura Virus forms cytoplasmic inclusion bodies through phase separation and proviral association of cellular protein Kinase R and stress granules protein TIA-1

**DOI:** 10.1101/2023.12.13.571494

**Authors:** Sharmistha Sarkar, Surajit Ganguly, Nirmal K. Ganguly, Debi. P. Sarkar, Nishi Raj Sharma

**Author notes:** **Abbreviation:** (Immunofluorescence assay; IFA, T-cell intracellular antigen; TIA-1, Poly A binding protein; PABP, Argonaute −2; Ago2, RasGTPase-activating protein-binding protein 1; G3BP1, eukaryotic initiation factor 3 eta; eIF3η, eukaryotic initiation factor 2 alpha; eIF2α, interferon inducible protein kinase R; PKR, Stress granule proteins; SGPs, Lipid droplets; LDs).

## Abstract

Negative-strand RNA viruses form cytoplasmic inclusion bodies (IBs) representing virus replication foci through phase separation or bio-molecular condensation of viral and cellular proteins, as a hallmark of their infection. Alternatively, mammalian cells form stalled-mRNA containing antiviral stress granules (SGs), as a consequence of phosphorylation of eukaryotic initiation factor 2α (eIF2α) through condensation of several RNA-binding proteins including TIA-1. Whether and how Chandipura virus (CHPV), an emerging human pathogen causing influenza-like illness, coma and death; forms IBs and evades antiviral SGs, remains unknown. By confocal imaging on CHPV-infected Vero-E6 cells, we found that CHPV infection doesn’t induce formation of distinct canonical SGs. Instead, CHPV proteins condense and co-localize together with SG-proteins to form heterogeneous IBs, which ensued independent of the activation of eIF2α and eIF2α Kinase, Protein Kinase R (PKR). Interestingly, siRNA-mediated depletion of PKR or TIA-1 significantly decreased viral transcription and virion production. Moreover, CHPV infection also caused condensation and recruitment of PKR to IBs. Compared to SGs, IBs exhibited significant rapidity in disassembly dynamics. Altogether, our study demonstrates that CHPV-replication co-optimizing with SG-proteins and revealing unprecedented proviral role of TIA-1/PKR, may have implication in understanding the mechanisms regulating CHPV-IB formation, and designing antiviral therapeutic.

**Importance:** CHPV is an emerging tropical pathogen reported to cause acute influenza-like-illness and encephalitis in children with very high mortality rate of ∼70%. Lack of a vaccines and an effective therapy against CHPV makes it a potent pathogen for causing an epidemic in tropical parts of globe. Given these forewarnings, it is of paramount importance that CHPV biology must be understood comprehensively. Targeting of host factors offers several advantages over targeting the viral components due to in general higher mutation rate in viral genome. In this study, we aimed at understanding the role of those cellular RNA binding proteins in CHPV replication, which form SGs. Our study helps understand participation of cellular factors in CHPV replication and could help develop effective therapeutics against the virus.

## Introduction

Being an emergent tropical pathogen causing acute fever and encephalitis among children, Chandipura Virus (CHPV) poses a huge threat to human health. First isolated in 1965 in Maharashtra State, India, from patients with febrile illness, CHPV, a member of the *Vesiculovirus* genus of the family *Rhabdoviridae* (1), is associated with severe human pathology which progresses rapidly from an influenza-like illness to coma and death (2). CHPV came to the limelight in 2003, when southern part of India experienced an outbreak of high mortality rate in which ∼350 children developed acute encephalitis and ∼200 died (2). The proposed carrier of the virus is the female *Phlebotomine* sandfly (2). CHPV has 150–165 nm long and 50–65 nm wide bullet-shaped morphology, as determined by transmission electron microscopy (3, 4).While a few CHPV outbreaks featuring a short incubation period and high mortality rate occurred in the past, their increased frequency in the recent years raise serious concerns about CHPV and necessitates preparedness against this deadly virus (2). CHPV genomic RNA is a single-stranded, negative sense (11,119 nucleotides, nts) and contains a 49 nt leader gene (l), five transcriptional units coding for viral polypeptides arranged in the order 3′ l-N-P-M-G-L-t 5′ separated by spacer regions and followed by a short non-transcribed 46 nt trailer sequence (t)(5). The complete genome sequence of CHPV was determined recently and comparative analysis of its deduced protein sequences showed CHPV to be phylogenetically distinct from its prototype *Vesiculovirus*, Vesicular Stomatitis Virus (VSV), but closely related to Isfahan virus (ISFV)(6).

Viruses are obligate parasites which hijack host cellular machinery for their multiplication, and subsequent transmission. However, the embedded vital intracellular machineries are not accessible to viruses at ease. To ensure their survival inside the host cells, in fact viruses essentially need to counter the multiple layers of intracellular resistance to replicate and establish their dominance for their propagation (7). One such layer in antiviral defense dimension in mammalian somatic cells comprise of the formation of two types of RNA granules, processing bodies (P-bodies, PB) and stress granules (SGs) (7–9). Although sharing a few components, both granules are physically and mechanistically distinct compartments with many unique biomarkers (10). While PB are ubiquitously formed during normal condition of cell growth and contain enzymes for RNA de-capping and degradation, (8, 11) and have been shown to store and degrade siRNA-or miRNA-guided mRNA (12, 13), SGs on the other hand are membraneless bodies produced to store mRNA during cell stress condition, lack de-capping/de-adenylating machinery and play a role in global translational arrest (8). The function of SGs is thus to serve as a central and dynamic storehouse to protect stored mRNA species and exchanged them with polysomes or PB for further translation or degradation, respectively (11, 14). The formation of SGs is in general, a defense mechanism against the stress to sequester mRNAs and arrest translation in order to save metabolic energy during stress condition. Besides containing several RNA-binding proteins [TIA-1, Ras-GAP (RasGTPase activating protein) SH3 domain-binding protein (G3BP1), Argonaute 2 (Ago2)], and mRNAs, SGs also contain 40S ribosomal subunits, and also many translation initiation factors including eIF3, eIF4G, eIF4E, and Poly(A)-binding protein PABP1(11, 15, 16). The exact mechanism of SGs formation is although not very clear but involves reversible condensation of mRNA and RNA binding proteins that stabilize or destabilize messenger RNA (mRNA).

In response to environmental stress, mammalian cells regulate translation-initiation as an efficient mechanism to conserve metabolic energy and nutrients which are abundantly consumed during protein synthesis. One of the central mechanisms in this regulation to arrest translational-initiation occurs by increasing the levels of phosphorylated eIF2α (p-eIF2α), which in turn leads to polysome disassembly triggering formation of SGs (17). Among various Kinases that phosphorylate eIF2α, host cell Protein kinase R (PKR) acts as a sensor for foreign or viral double stranded RNA (dsRNA) and in the process gets activated to induce eIF2α phosphorylation leading to biomolecular condensation of RNA and proteins for SGs formation (10, 18). This in turn, shuts down host cell translation and triggers the host cell antiviral response (19, 20). However, the viruses have evolved multiple strategies to bypass this response and even have been shown to utilize alternative ways of translation for their propagation(21). Several viruses show ability to suppress the formation of SGs (22–26). For instance, Poliovirus C3 protease and Semliki Forest virus nsP3 target G3BP, which regulates initial phase of SGs assembly (24, 27–29). Influenza virus NS1 inhibits the activity of dsRNA-activated PKR(22, 23). Interestingly, Hepatitis C Virus co-opts SGs and thus induces SGs formation for its replication and production (30, 31).

Several reports suggest that low complexity sequence domain mediate the liquid–liquid phase separation in RNA binding proteins which contribute to the formation of SGs or membrane-less bio-molecular condensates (32, 33).In the context of viruses, bio-molecular condensation seems to provide viruses with numerous advantages that allow them to establish replication organelles, assemble viral particles, and even evade innate immune responses (34). It has been recently demonstrated that phase separation of the human adenovirus intrinsically disordered 52-kDa protein plays a critical role in the coordinated assembly of infectious progeny particles, and for the organization of viral structural proteins into biomolecular condensates (35).

Recent studies suggested that like SGs or PBs, viral IBs also have properties of bio-molecular condensates (36).We recently reported a varying degree of disorder in all five CHPV proteins, with the maximum level of intrinsic disorder propensity found in Phosphoprotein (P) (5). In pursuit of the identification of host factors involved in CHPV replication, we investigated three key aspects of CHPV infection. These included 1) understanding the propensity of CHPV proteins to undergo condensation and forming CHPV replication factories or inclusion bodies or IBs 2) participation of cellular proteins in these factories and 3) CHPV evasion of the antiviral SGs formation. Here, by Immunofluorescence assay (IFA) on CHPV infected cells for nucleocapsid (N) and Large (L) protein detection, we found that CHPV proteins indeed show condensation and co-localize to form a heterogeneous population of puncta. Although, CHPV infection did not induce distinct SGs but it promoted condensation of multiple SG markers (TIA-1, G3BP1, PABP, Ago2, and eIF3η) which showed co-localization with large CHPV punctas referred as CHPV-IBs. Interestingly, recruitment of SG proteins to IBs ensued independent of PKR/eIF2α phosphorylation. Nevertheless, CHPV infection caused condensation and recruitment of PKR to IBs while siRNA-mediated depletion of PKR or TIA-1significantly decreased virion production. The IBs exhibited cycloheximide sensitivity and in comparison to SGs, significant haste in disassembly dynamics. Taken together, our study demonstrates TIA-1 and PKR as proviral factors in CHPV replication. This may have implication in understanding the mechanisms regulating CHPV-IB formation, and designing anti-viral therapeutics. Moreover, this study not only provides the first insight about the cytoplasmic viral IBs formation during CHPV infection but also pave the way for a plethora of more investigations to understand the mechanism of CHPV-IBs formation during its replication.

## Results

### CHPV infection induces condensation of viral nucleocapsid (N) and Large (L) proteins to form cytoplasmic IBs

We initiated this study by monitoring cytopathic changes in live CHPV-infected Vero-E6 cells. Here, we observed a rapid change in the morphology of infected cells (CHPV MOI= 5), turning from elongated to round in as early as 4 hrs P.I. in ∼5% of cells; which by 7 hrs P.I., could be detected in ∼90 % (Fig S1A-B). CHPV infection is known to induce apoptosis in a variety of mammalian host cells, although to a varying degree (37, 38).To understand a correlation between roundening and apoptosis in CHPV infected Vero cells, we employed a commonly used tetrazolium dye based MTT assay to assess the cellular metabolic activity. Here, the time course data of escalating roundening of cell correlated with that of cell death and suggested roundening of cells precedes apoptosis in infected cells (Fig S1B).

Next, to understand the propensity of CHPV proteins to undergo condensation during the course of infection, we performed IFA on CHPV infected cells, to detect expression and intracellular localization of two CHPV proteins (Nucleocapsid ‘N’ and Large ‘L’) (Fig 1A). While, N encapsidates the genome in a ratio of one N per nine ribonucleotides, protecting it from nucleases, L protein is a RNA-directed RNA polymerase that catalyzes the transcription of viral mRNAs, their capping and polyadenylation. Notably, N represents the first and most abundant, while L as the largest but the least abundant protein, to express during CHPV infection. Here, by confocal microscopy, we observed visible appearance of both proteins after 3hrs P.I., and expression of N protein much more abundant than L protein, as expected (Fig 1B). Cytosolic heterogeneous expression of N protein was observed diffused as well as localized in puncta of variable size. However, we observed maximum percentage of cells displaying the highest average number of CHPV-N puncta at 4 hrs P.I., with ∼5-10% cells forming >50 puncta (Fig S1C, Fig 1C).

**Fig.1.**
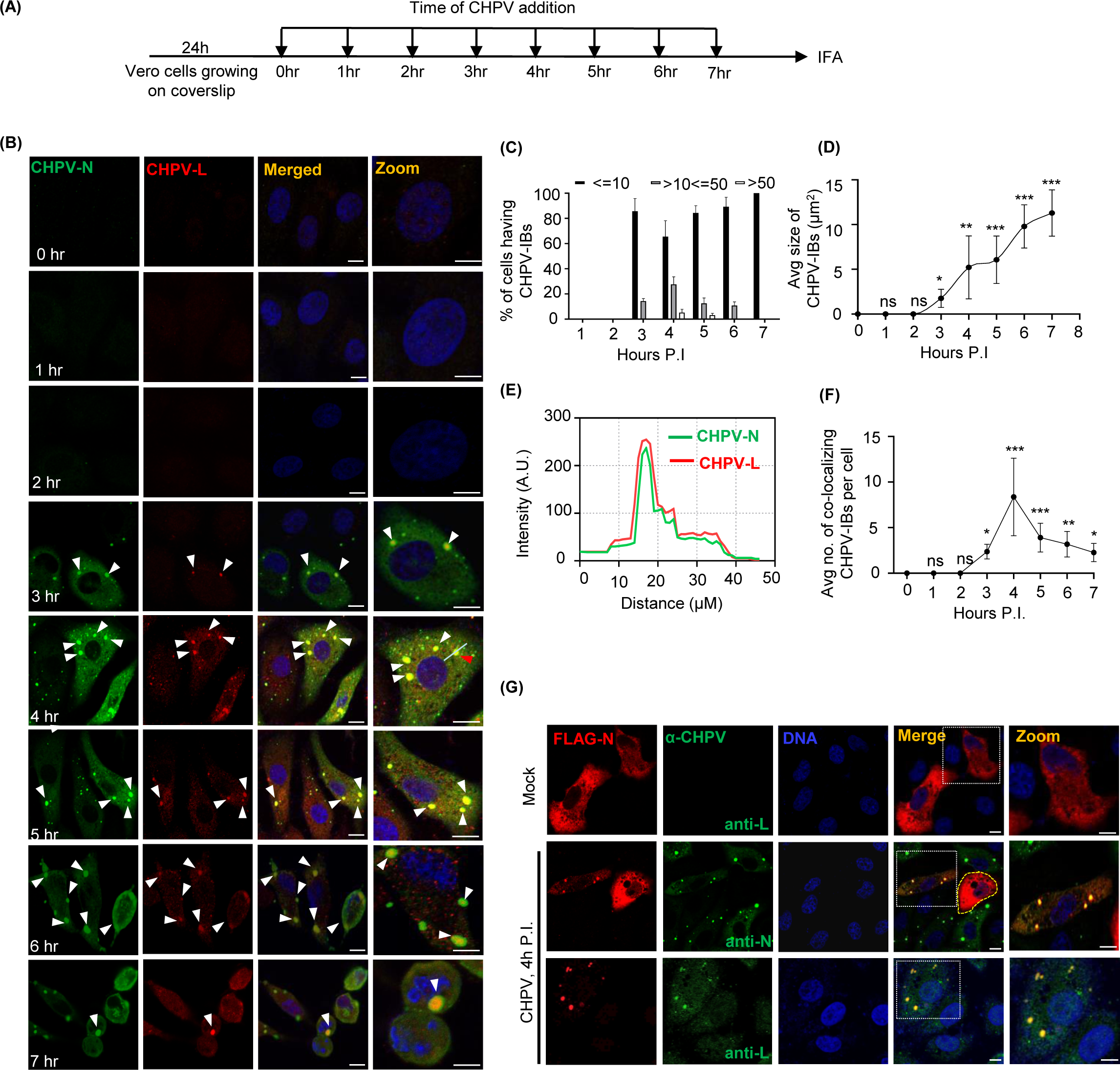
CHPV proteins condense and co-localize together to form cytoplasmic IBs in infected cells. **(A)** Outline of experimental design for the time course of CHPV infection in Vero cells grown on cover slips at the indicated time and infected cells were collected at the same time for further fixation and immunostaining. **(B)** Time course of CHPV infection. Vero Cells were infected either with 5 MOI live CHPV for indicated time or with heat-inactivated (56°C/20 min) CHPV (HI-CHPV) for 7 hrs. Cells were fixed and co-immunostained for CHPV-N and CHPV-L using specific antibodies. The nuclei were counterstained with Hoechst dye. Scale bar = 10 µm. **(C)** Bar graph showing percentage of cells in (B) with less than or equal to ten (<=10), more than ten but less than or equal to fifty (>10<=50) or more than fifty (>50) CHPV-N puncta at each time point. The error bars represent SD from three independent experiments. **P<0.01 in Chi-squared test. **(D)** Graphical representation of the average size of CHPV-N punctas over the course of infection CHPV-N puncta from at least 25 cells from each time point of infection were measured using ImageJ. The error bars represent SD from three independent experiments. **P<0.01 in Chi-squared test. **(E)** The distribution of signal intensities for CHPV-N and CHPV-L over selected cells from panel B (red arrow) using the Image J software. AU, arbitrary units. **(F)** Graph representing average no. of co-localizing CHPV-IBs per cell over the time course of CHPV infection calculated using Image J (n>35 cells). **(G)** CHPV infection dependent condensation and co-localization of CHPV proteins. Vero cells were transfected with pFlagCMV6a-CHPV-N and 24 hrs later infected with 5MOI CHPV. The cells were further co-immunostained for the detection of flag-tagged CHPV-N (using anti-Flag) in combination with anti-N antibody or anti-L antibody for detection of N and L protein expressed during CHPV infection. Uninfected or CHPV infected cells are separated by dashed white borderlines. Uninfected cell outlined by yellow dashed line. The nuclei were counterstained with Hoechst dye. Scale bar = 10 µm.

L protein, albeit expressed exclusively in the form of puncta, colocalized with large punctas of N (Fig. 1B,1D-E, S1D), suggesting these to be the CHPV inclusion bodies, herein referred as “CHPV-IBs” or “IBs”. The phenomenon of CHPV-IBs formation in Vero cells appeared to be diverse ranging from maximum 5-10 IBs/cell at 4hr (Fig. 1B & 1F), which gradually decreased to as low as 1-2 IBs/cell at 7 hrs P.I., as small IBs coalesce to form large IBs by the end of 7 hrs, resulting in a steady increase in average size of IBs (Fig 1D).

In order to examine independent ability of N-protein to form puncta, we overexpressed a recombinant CHPV-N protein, fused to Flag tag at its N-terminus, in Vero cells and confirmed its size and expression by immunoblotting technique using anti-Flag antibodies (Fig S1C). IFA on Vero cells ectopically expressing N protein, in the absence of CHPV infection, showed its diffused distribution in the cytoplasm (Fig 1G, uppermost panel). CHPV infection in these cells however dramatically changed this distribution to puncta which showed colocalization with both virally encoded CHPV-N and CHPV-L proteins (Fig 1G, middle and lower panel). Taken together, these results indicated that CHPV proteins (N and L) condense and colocalize to form cytoplasmic IBs during CHPV infection.

### Recruitment of multiple stress granules proteins (SGPs) to CHPV-IBs

Virus infection, in general, causes stress on multiple biosynthetic pathways in host cells (39). In response, mammalian somatic cells undergo global translational arrest and in consequence, produce SGs to store stalled mRNA and prevent viral replication (8, 40). To understand whether CHPV infection induce SGs and their interplay with CHPV-IBs, we performed IFA on infected cells for several SGPs, as shown in the Fig. 2A. Among these, TIA-1, an RNA-binding protein, recognize the translationally arrested mRNPs and nucleates the assembly of SG through its prion-like aggregation property (41). As expected, in absence of CHPV infection, we observed predominant nuclear staining of TIA-1, but no visible distinct SGs (Fig. 2B, upper panel). In contrast, CHPV infected cells displayed TIA-1 forming 5-10 cytoplasmic puncta (Fig. 1B lower panel). Since all TIA-1+ puncta also showed co-localization with IBs represented by CHPV-N immunostaining (Fig. 2B, C), it was suggested that although CHPV infection does not induce formation of distinct canonical TIA-1+ SGs, but somehow induce TIA-1 localization to CHPV-IBs. To address whether more SGPs are recruited to IBs, next we performed IFA to determine IBs association with three endogenous (PABP1, Ago2, eIF3η) and one ectopically expressed (GFP-tagged G3BP1)marker of SGs. Here, Ago2 or Argonaute2 is a core component of RNA induced silencing complex (RISC) to play its key role of catalytic engine that drives mRNA cleavage in RNA interference (42). However, cellular stress leads to rearranged and increased association of Ago2 with the coding and 3′ UTRs of mRNAs and its recruitment to SG(43, 44). Ras-GTPase-activating protein (GAP)-binding protein 1 (G3BP1) is a multi-functional RNA binding protein, best known for its role in triggering the assembly and dynamics of SGs (10, 45). Likewise, eIF3η, a subunit of eIF3 complex to control the assembly of the 40S ribosomal subunit and stabilizing eIF2-GTP-Met-tRNAiMet complex association, is also an authentic marker of SGs (46). Interestingly, immunostaining of all the RNA binding proteins (PABP1, eIF3η, and Ago2), in contrast to their cytoplasmic diffused distribution in uninfected cells, exhibited puncta and their colocalization with CHPV-IBs (Fig. 2D-E, 2F-G, 2H-I). Nevertheless, ectopic expression of GFP-tagged G3BP1 was observed in uninfected and CHPV infected cells. When compared to its expected diffused distribution in uninfected cells, we found that G3BP1 also undergoes condensation and forms puncta in infected cells (Fig. 2J, compare upper and lower panel). CHPV-N co-localization and TIA-1+ staining of these G3BP-1 puncta confirmed (Fig. 2J & K) them as CHPV-IBs. In contrast to GFP-G3BP-1, GFP alone, which served as a control protein, did not form visible puncta and also did not colocalize with IBs positive for TIA-1 and CHPV-N (Fig. S2A&B).

**Fig. 2.**
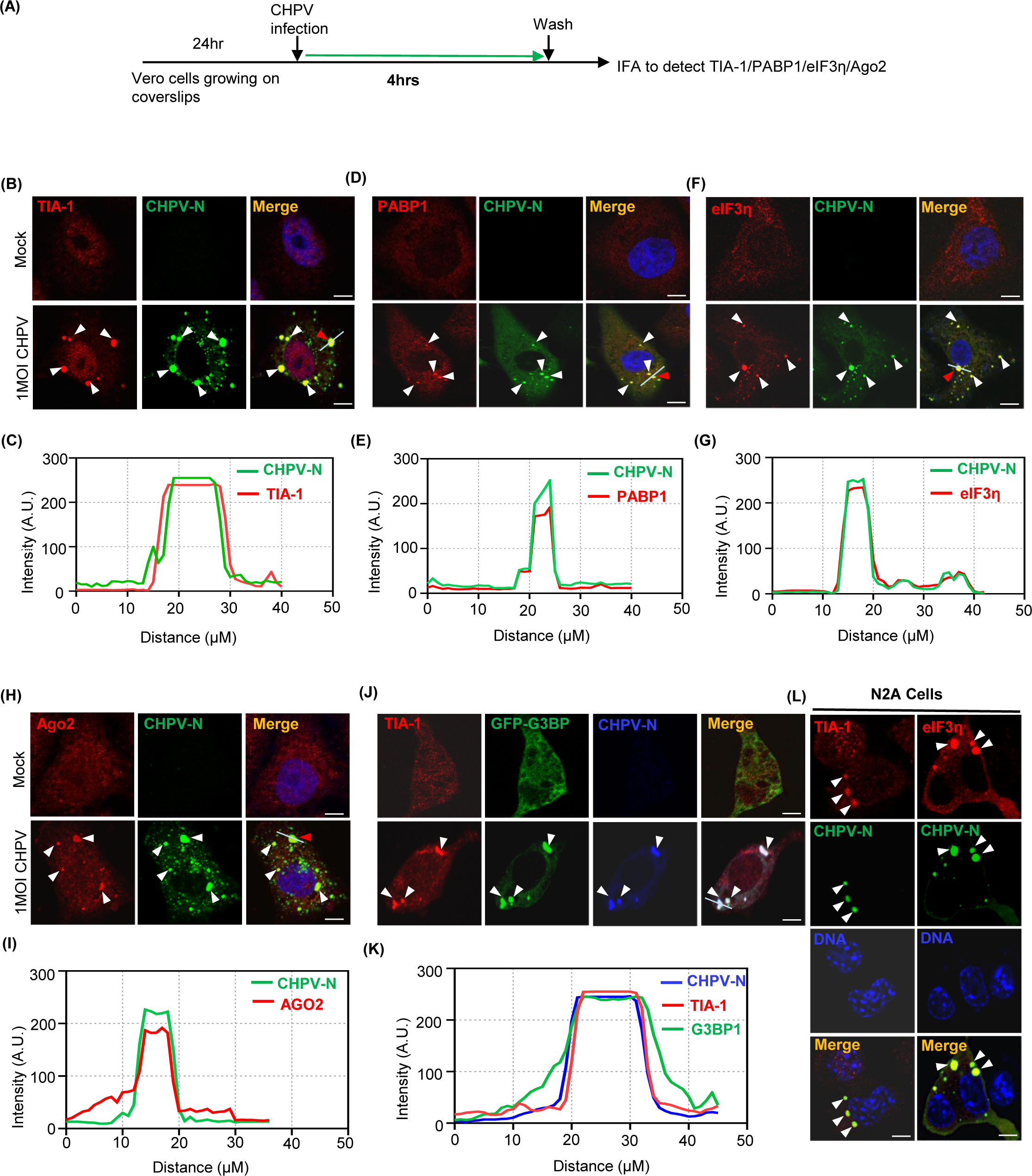
Multiple cellular SGPs associate with CHPV-IBs. **(A)** Outline of the experimental design for detection of SGPs in CHPV infected cells. Vero cells grown on cover slips were infected with 1 MOI CHPV for 4 hrs and then processed for IFA to detect CHPV-N in combination with SGPs using specific antibodies. **(B-I)** CHPV-IBs co-localize with endogenous SGPs (TIA-1, PABP1, eIF3η, and Ago2). Vero cells were infected with 1 MOI CHPV for 4 hrs. Cells were then fixed with 4% PFA and immunostained for detection of endogenous markers. TIA-1 (red) in (B), PABP1 (red) in (D), eIF3η (red) in (F) and Ago2 (red) in (H) were detected along with CHPV-N (green) using specific antibodies. Graphical representations in (C, E, G, I) shows the distribution of signal intensities across the line for each SGP and CHPV-N over selected cells from panel B, D, F and H (red arrows) using the Image J software. AU, arbitrary units. The nuclei were counterstained with Hoechst stain. Scale bar = 10 µm. **(J-K)** Ectopic expression and co-localization of GFP-G3BP1 with CHPV-IBs. Vero cells transfected with a GFP-G3BP1 or GFP (Fig S2A) alone expressing vector for 24 hrs were infected with 5MOI CHPV for 4 hrs. The cells were stained for TIA-1 (Red) and CHPV-N (Blue) using their respective antibodies. Graphical representations in (K) shows the distribution of signal intensities across the line for GFP-G3BP1, TIA-1 and CHPV-N over selected cell from panel J (red arrows) using the ImageJ software. AU, arbitrary units. Scale bar = 10 µm. **(L)** CHPV-IBs co-localize with endogenous SGPs in N2A cells. N2A cells were infected with 5 MOI CHPV for 4 hrs. The cells were then fixed with 4% PFA and immunostained for TIA-1(red), eIF3η (red) and CHPV-N (green) using their respective antibodies. The nuclei were counterstained with Hoechst stain. Scale bar = 10 µm.

Apart from SGs, mammalian cells also form Lipid droplets or LDs, as storage organelles at the center of lipid and energy homeostasis. They have a unique architecture consisting of a hydrophobic core of neutral lipids, which is enclosed by a phospholipid monolayer that is decorated by a specific set of proteins. LDs have shown to be highly dynamic, ubiquitous organelles, which are found in virtually all types of cells from prokaryotes to eukaryotes. They consist mainly of triglycerides and sterol esters, but also harbor other lipid species such as diacylglycerols, retinyl esters and ceramides (47, 48), and can also activate the synthesis of bioactive lipid mediators (49). LD biogenesis has been rapidly detected after infection with a number of different viral and non-viral pathogens. Next, to understand any physical association between IBs and lipid droplets (LD), we performed IFA using a BODIPY lipid droplet dye. This lipophilic dye can pass through the cell membrane and distribute within the cell allowing it to specifically label cellular lipid droplets. Though, we did not find colocalization of CHPV-IBs with LDs, there was a drastic drop in the number of LDs suggesting that this effect to be associated with increased toxicity in the infected cells. Taken together, it appeared that IBs do not associate with LDs (Fig. S2C&D).

CHPV exhibits neurological manifestation in young children and can induce neurological symptoms in suckling mice, thus indicating its neurotropic characteristics (50). This led us to confirm CHPV neurospecificity for IBs formation and their association with SG proteins in Neuro 2A (N2A) cells. Interestingly, similar to Vero Cells, N2A cells displayed formation of ∼3-5 CHPV-N IBs/cell which showed co-localization with condensed puncta of TIA-1 and eIF3η in cytoplasm (Fig. 2L). In the absence of CHPV-N puncta in uninfected N2A cells, TIA-1 and eIF3η did not undergo visible condensation (Fig. S2E). Collectively, these results indicate that the ability of CHPV infection to trigger condensation and recruitment of several cellular SGPs to form Ibs is cell type independent.

### CHPV-IBs and canonical SGs are distinct in terms of disassembly dynamics

Cycloheximide (CHX), a polysome stabilizing drug, is a translational inhibitor that stalls the elongation of translation without allowing the disassembly of the polysomes and thus prevents the formation and/or maintenance of SGs (51, 52). Since viral replication exclusively relies on cellular translational machinery, we anticipated CHX to be a potential inhibitor of IBs and thought to use this ability to understand and compare the dynamics of assembly/disassembly of IBs with that of canonical sodium arsenite (SA, NaAsO₂) induced SGs. In agreement with previous observations (14, 53), SGs induction presence or absence of CHX, and monitored by CHPV-N and PABP-1 immunostaining followed by confocal microscopy. Here, in the absence of CHX, SG s disappeared at a very slow rate (∼100 min for 50% reduction) (Fig. 3F-G). However, in the presence of CHX, the rate of their disassembly became two times faster (∼50 min for 50% reduction) (Fig. 3E-G). In comparison, CHPV-IBs, the disassembly of which could only be imposed in the presence of CHX, interestingly exhibited a relatively faster rate of disassembly than SGs (∼30 min for 50% reduction) (Fig. 3F-G). Altogether, this implied not only towards the IBs dependence on viral mRNA translation for their induction and maintenance, but also their higher sensitivity to CHX during the disassembly process, thus differentiating from the canonical SGs.

**Fig. 3.**
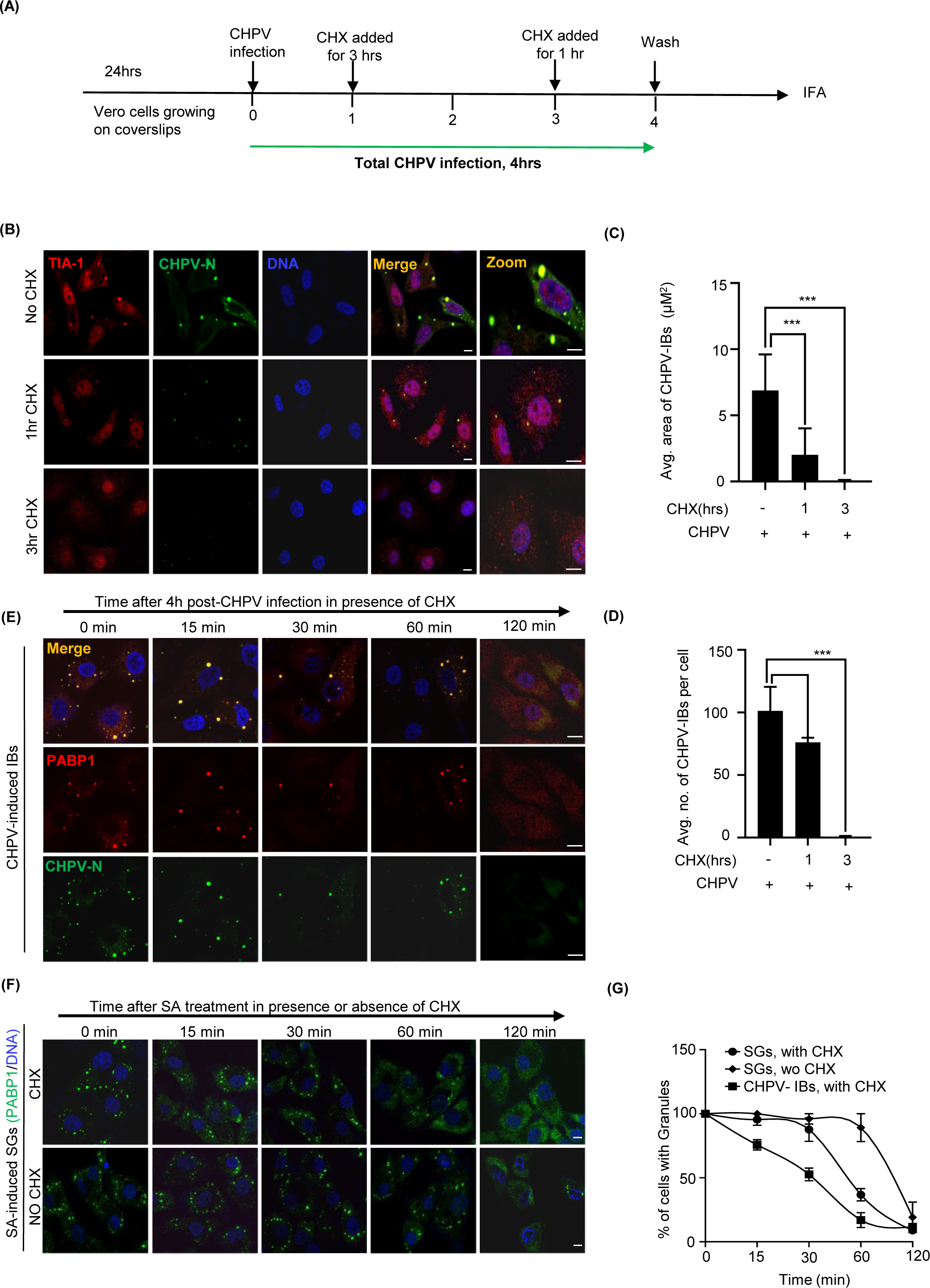
Cycloheximide (CHX) treatment reduces the size and number of CHPV-IBs. **(A)** The schematic shows outline of experimental design. Vero cells grown on coverslips were infected with CHPV 5MOI for 4 hrs. CHX was added to cells at the indicated times for 3 hrs and 1 hr treatment respectively. **(B-C)** CHPV-IBs are sensitive to CHX treatment. Vero cells were infected with CHPV 5MOI either in the absence or presence of CHX (50 μg/ml) for 1 hr and 3 hrs as shown in (A). After completion of 4 hrs of infection, cells were fixed with 4% PFA and immunostained for TIA-1 (red) and CHPV-N (green). The nuclei were counterstained with Hoechst dye. Scale bar =10 µm. **(C-D)** graphical representation of the number (C) and size (D) of CHPV-IBs in an effect of CHX treatment as mentioned in (A-B). **(E-F)** Comparison of Kinetics of CHPV-IBs and SGs disassembly upon CHX treatment. Vero cells were either infected with 5MOI CHPV for 4 hrs (E) to induce IBs or treated with 1mM SA for 40 min to induce SGs (F). In both (E) and (F), cells were either left untreated or subsequently treated with CHX (50μg/ml) for 15 min, 30 min, 1 hr and 2 hrs to compare the dynamics of SGs and IBs disassembly. The cells were then fixed and co-immunostained for CHPV-N and PABP1. The nuclei were counterstained with Hoechst dye. Scale bar = 10 µm. **(G)** Graph showing comparison between the CHPV-IBs and canonical SGs with respect to their kinetics of disassembly in the presence of CHX. Untreated SGs without CHX served as a control.

### TIA-1 and PKR play a pro-viral role in CHPV replication

So far, we found that CHPV infection induce condensation of TIA-1 and other SG-proteins and association with IBs. To understand whether this association of TIA-1 is pro-viral or anti-viral, we examined CHPV virion production in Vero-E6 cells with or without siRNA-mediated TIA-1 knockdown. Although, production of interferons is defective in Vero cells (54) which allows efficient viral replication, yet among its several effector molecules, double-stranded RNA (dsRNA)-dependent protein kinase (PKR), an enzyme with multiple effects in cells, is capable of playing a critical role in the antiviral defense mechanism of the host with important biological functions, including translational regulation (55). Thus, in parallel, we also performed PKR knockdown to validate the hypothesis for antiviral role of PKR in CHPV infection. By using a TIA-1 or PKR specific siRNAs, the dose of siRNA for a significant knockdown was optimized to achieve almost ∼70% and ∼85% of TIA-1 and PKR expression respectively (Fig. S4A-D). Next, we divided siRNA-transfected Vero cells into four groups to assess the effect of TIA-1 /PKR depletion on CHPV gene expression and its replication as shown in Fig. 4A. Cells were infected with 1MOI for 4 or 6 hrs as shown in Fig. 4A, for CHPV-N to visualize using or detect its mRNA or protein level. Alternatively, cells were infected with low MOI (0.01) CHPV for 18 hrs, followed by collecting supernatant for virion quantification using plaque assay. Here, we found that efficient knockdown of TIA-1 or PKR expression from Vero cells resulted in significantly decreased production of not only CHPV-N protein (Fig. 4B-C & Fig. 4D-E) as detected by western blotting; but also of CHPV-N mRNA (Fig. 4F) quantified by Real-time PCR. It may be noted that silencing PKR expression did not change total levels of either eIF2α or its phosphorylated form, suggesting this effect to be specific to CHPV-N protein (Fig. 4B-C). Moreover, IFA results corroborated with that of western blotting showing that siRNA knockdown of PKR or TIA1 expression in Vero cells led to ∼70-80% decrease in CHPV-N expression over the cells with the normal level of PKR or TIA-1 expression (Fig. 4D-E and Fig.S4E-F). Nevertheless, silencing of TIA-1 or PKR expression also led to a concomitant reduction of infectious virions in Vero culture supernatants (Fig.4G-H). Next, to test whether antagonistic effect of PKR or TIA-1depletion on CHPV growth resulted in increased cell viability, we performed phase contrast microscopy and MTT assay to check cell viability. Here, we found decreased roundening and increased cell viability of CHPV infected cells after TIA-1 or PKR knockdown (Fig. 4I). Altogether, this data suggested TIA-1 and PKR playing a proviral role in the CHPV replication.

**Fig. 4.**
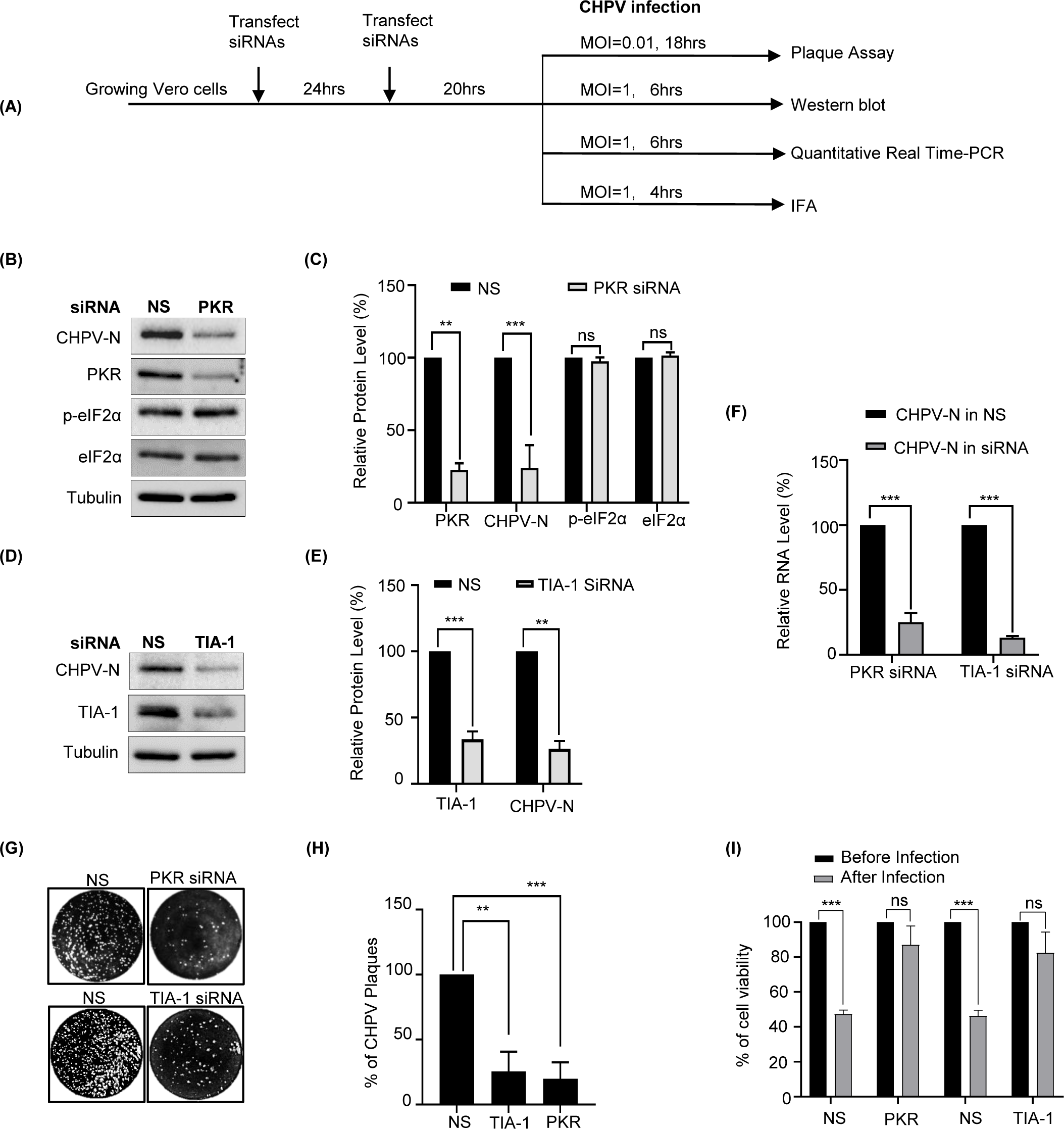
Silencing of TIA-1 or PKR expression decreases CHPV production. **(A)** Outline of experimental design for siRNA transfection following CHPV infection in Vero cells. After siRNA transfections (twice after an interval of 24 hrs), cells were equally divided into 4 sets. Each set was infected with the indicated MOI of CHPV for the indicated time for further detection of virions by plaque assay, quantification of viral RNA by qPCR and viral protein by western blotting or IFA. **(B)** Western blot detection of PKR, CHPV-N, p-eIF2α, eIF2α and tubulin in CHPV infected (1 MOI, 6 hrs) Vero cells after siRNA mediated silencing of PKR expression. **(C)** Graphical representation of the relative amount of PKR, CHPV-N, p-eIF2α, and eIF2α. The relative intensity of each protein band in PKR-siRNA sample, after normalizing to tubulin, was calculated over that of the NS-siRNA control. The error bar indicates mean ±SD (n=3). **(D)** Western blot detection of TIA-1, CHPV-N and tubulin in CHPV infected (1 MOI, 6 hrs) Vero cells after siRNA mediated silencing of TIA-1 expression. **(E)** Graphical representation of the relative amount of TIA-1 and CHPV-N. The relative intensity of each protein band in TIA-1-siRNA sample, after normalizing to tubulin, was calculated over that of the NS-siRNA control. The error bar indicates mean ±SD (n=3). **(F)** Graphical representation of the relative amount of CHPV-N mRNA in CHPV infected (1MOI, 6 hrs) Vero cells after siRNA mediated silencing of PKR/TIA-1 as compared to that in NS-siRNA control. The error bars indicate mean ±SD (n=3). **(G)** Analysis of CHPV virion production after siRNA knockdown of PKR or TIA-1. As shown in (A), siRNA transfected Vero cells were infected with CHPV (0.01 MOI) and allowed to replicate CHPV for 18 hrs. Cell culture supernatants obtained from the infected cells were used for plaque assay as described in Material and methods. **(H)** Graphical representation of the quantification of plaques obtained in (G). Cells were fixed and stained with crystal violet at 18 hrs P.I., as described in Material and methods. The error bar indicates mean ±SD (n=3). **(I)** Graphical representation of the quantification of cell viability before/after CHPV infection (1MOI, 6hrs) in Vero cells after siRNA mediated silencing of PKR/TIA-1 as compared to that in NS-siRNA control. The error bars indicate mean ±SD (n=3).

### PKR undergoes condensation and associates with CHPV-IBs

TIA-1 co-localization with CHPV-IBs made us to speculate a unique proviral role of PKR in CHPV replication and its participation in the formation of CHPV-IBs. To work on our hypothesis, we performed IFA on CHPV infected Vero cells. In addition, to address whether this is a unique association of PKR with CHPV-IBs, we also sought to compare between IBs and canonical SGs. In this pursuit, we also treated one set of cells with 1mM SA for 40 mins to induce SGs. Both IBs and SGs were co-immunostained with anti-PKR antibodies. Additionally, SGs were also co-immunostained with anti-PABP1 antibodies together with anti-TIA-1 to compare and confirm the architecture of SGs. Here, in normal Vero cells, while PABP1 exhibited exclusively cytoplasmic immunostaining, both TIA-1 and PKR showed nucleocytoplasmic distribution (Fig. S5A). SA induced SGs stained positive for TIA-1 and PABP1 but stained negative for PKR (Fig. 5A). The graphical representation of the measured signal intensity along a line across the SGs showed colocalization of TIA-1 and PABP1, but not of PKR (Fig. 5B). On the other hand, CHPV-IBs, represented by CHPV-N immunostaining, also stained positive and displayed colocalization with PKR which as compared to uninfected cells, showed condensation in infected cells (Fig. 5C-D). Taken together, the data suggested that similar to TIA-1, PKR also associate with IBs to play a proviral role in CHPV replication.

**Fig. 5.**
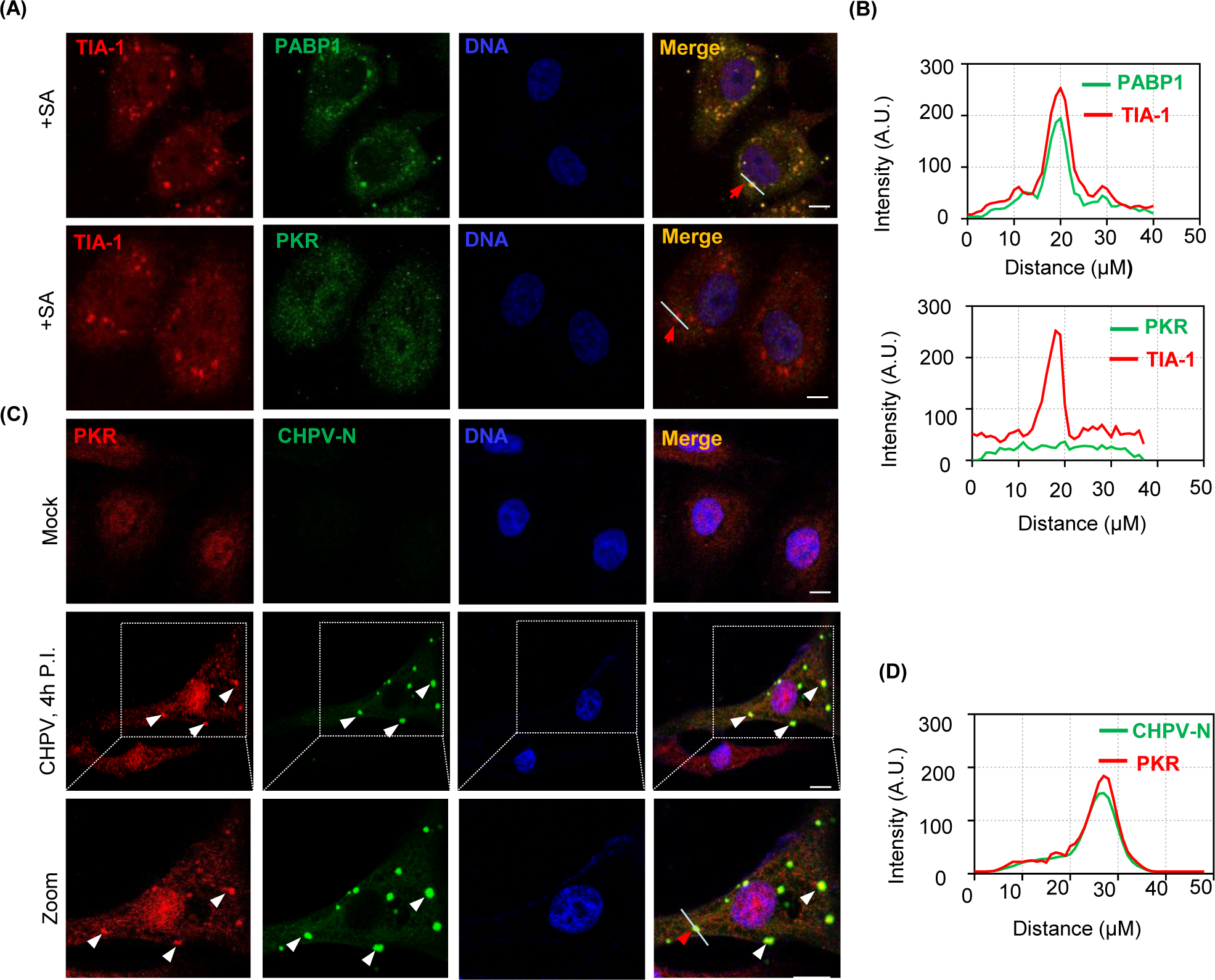
PKR associates with CHPV-IBs but not with SGs. **(A)** Vero Cells were treated with 1mM SA for 40 min to induce SGs. Later cells were fixed with 4%PFA and processed for IFA to detect TIA-1/ PABP1 or TIA-1/PKR for SGs. The nuclei were counterstained with Hoechst dye. Scale bar = 10 µm. **(B)** Graphical representations of the intensity plot across the line drawn in the images (Fig A: upper and lower panel of SGs) for individual measurements in arbitrary units (AU) is shown after normalization to maximum intensities using Image J software. **(C)** Vero Cells were infected by CHPV 1MOI for 4 hrs to induce IBs. Cells were fixed with 4% PFA and then processed for IFA to detect CHPV-N/ PKR for CHPV-IBs. The nuclei were counterstained with Hoechst dye. Scale bar = 10 µm. **(D)** Graphical representations of the intensity plot across the line drawn in the images (Fig C: lower panel of CHPV-IBs) for individual measurements in arbitrary units (AU) is shown after normalization to maximum intensities using Image J software.

### CHPV-IBs form independent of PKR and eIF2α phosphorylation

Phosphorylation of α subunit of eIF2α stalls mRNA translation and promotes condensation and aggregation of TIA-1 to form the SGs for storing mRNA (56). Of the four Kinases which phosphorylate eIF2α (GCN2, general control non-derepressible 2; PKR, Protein kinase R; PERK, PKR-like endoplasmic reticulum kinase; and HRI, heme regulated inhibitory kinase), PKR can be phosphorylated by viral infection (20, 57) and alternatively by SA (44, 58). While Virus infection can generate dsRNA and activates PKR through binding of dsRNA to the dsRNA-binding domain (RBD) of PKR, SA activates PACT to bind and activate PKR (58, 59) (Fig. 6A). PKR possess at least 15 autophosphorylation sites, however phosphorylation at Thr 446 and Thr 451 is critical for its activation, and subsequent phosphorylation of eIF2α (60, 61). We hypothesized that, similar to SGs, condensation of SGPs and their recruitment to IBs during CHPV infection may be dependent on phosphorylation of eIF2α and also assumed PKR responsible for this function possibly due to its association with IBs. Accordingly, we examined whether CHPV infection could affect PKR/ eIF2α phosphorylation. To correlate kinetic production of CHPV proteins with both total eIF2α and phosphorylated eIF2α, we infected Vero cells with two doses of CHPV (1 MOI and 5 MOI) for the indicated time as shown in (Fig. 6B and S6A). As mentioned in Fig. 5B, we also treated uninfected Vero cells with 1mM SA for 40 min before collecting cell lysates for Western blotting analysis, to serve as a positive control for PKR and eIF2α phosphorylation. In this assay, while SA treatment induced phosphorylation of both PKR and eIF2α, CHPV infection did not induce such activity over that of uninfected cells [(Fig.6B, lane 7 vs lanes 1-6), and Fig. 6C]. It may be noted however that reprobing the same membrane with specific antibodies showed activation of p38 phosphorylation by CHPV infection, as reported earlier (62), with the total level of p38 protein remaining the same.

**Fig. 6.**
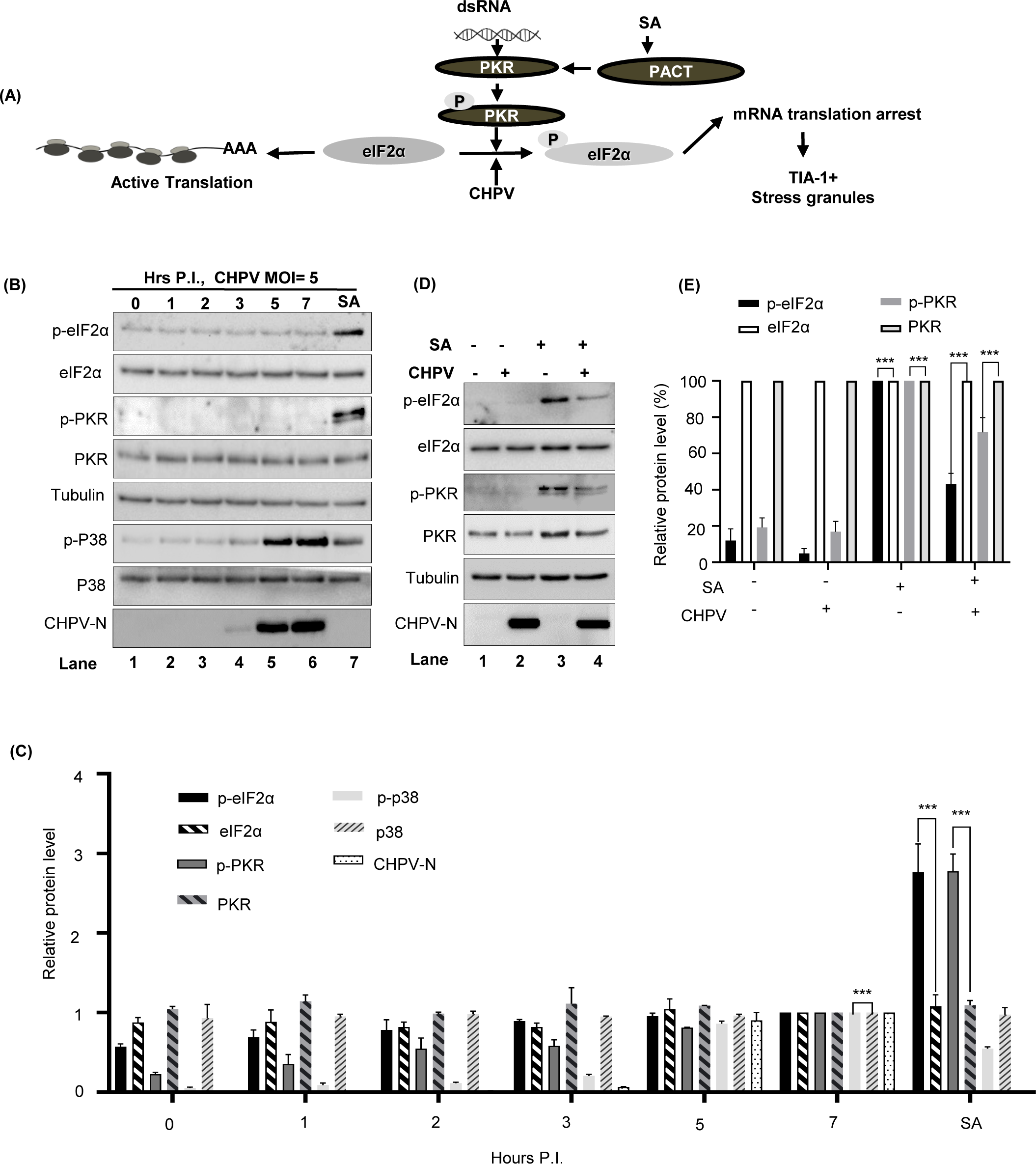
CHPV infection induces p38 phosphorylation but does not phosphorylate PKR/eIF2α. **(A)** Schematic diagram showing the cellular pathway p-PKR/p-eIF2α/SG governing SG formation. PKR direct activation by dsRNA or indirectly by SA through PACT is shown. **(B)** Kinetics of p38 activation with no change in p-PKR/p-eIF2α. Vero cells were infected with CHPV (5 MOI) for the indicated time. The phosphorylated forms of eIF2α (p-eIF2α, Ser51), PKR (p-PKR) and p38 (p-p38) were measured by Western blot analysis using phospho-specific antibodies. Total level of these kinases was determined by a pan-antibody. Tubulin served as a loading control. **(C)** Graphical representation of the relative amount of p-eIF2α, eIF2α, p-PKR, PKR, p-p38 and p38 protein in each sample after normalizing to tubulin was plotted over the time when the sample was collected (6B) with the protein level in lane 6 (7 hrs of CHPV infection) set as 1. The error bar indicates mean ±SD (n=3). **(D-E)** CHPV infection inhibits SA induced eIF2α phosphorylation. Vero cells with or without CHPV infection were left untreated or treated with 0.5mM SA for 30 min before cell lysate preparation for Western blotting with corresponding antibodies. CHPV-N was blotted as an indication for viral infection. (E) Relative amount of p-eIF2α, totaleIF2α, p-PKR, and total PKR in each sample after normalizing to tubulin was measured and plotted in bar graphs for comparison, with each protein level in lane 3 being set to 100%. The error bar indicates mean ±SD (n=3).

Next, we also wanted to understand the effect of CHPV infection on SA induced PKR/eIF2α phosphorylation. For this, Vero cells either uninfected or infected with CHPV, were treated with 1mM SA for 40 min and were analyzed by western blotting for phosphorylation status of both eIF2α (p-eIF2α) and PKR (p-PKR). As expected, SA was found to induce eIF2α phosphorylation ∼10 times greater than the basal level (Fig.6D lane 3 vs lane 1). In contrast, CHPV infected cells exhibited remarkable inhibition (∼60%) of eIF2α phosphorylation upon SA induction (Fig.6D lane 4 vs lane 3, Fig.6E), with no detectable changes in the total eIF2α protein signal. Moreover, similar effect of CHPV infection on partial inhibition of PKR phosphorylation (∼30%) was also observed without any effect on total PKR levels (Fig. 6D lane 4 vs lane 3, Fig. 6E). This data indicated that CHPV infection is partially inhibitory for PKR and eIF2α phosphorylation. Taken together, the data suggested that condensation and recruitment of SGPs to CHPV-IBs is independent of PKR/eIF2α phosphorylation.

## Discussion

Despite its huge potential to cause a deadly infection, how CHPV utilizes or hijacks cellular factors to replicate inside the cells, remains largely unknown. Negative-strand RNA viruses, in general, induce the formation of cytoplasmic Inclusion bodies (IBs) which involves phase separation of viral and cellular proteins. Numerous reports recently came out to suggest viral proteome contain intrinsically disordered proteins (IDP) and IDP regions (IDPR), which are proteins or protein regions that lack unique (or ordered) three-dimensional structures (5, 63, 64). Of late, a number of reports also suggested involvement of IDP and IDPR in cell signaling (65), assembly of cellular SGs (66, 67) and nonetheless, in condensation and aggregation of viral and cellular proteins to form IBs (68). In this direction, we published a report emphasizing on the varying degree of disorder in all five CHPV proteins, with the maximum level of intrinsic disorder propensity found in Phosphoprotein (P) (5). To further investigate the propensity of CHPV proteins for phase separation and for exploration of its host factor usage, a Vero cell line, as reported earlier (38), was investigated as a suitable host system for CHPV replication. While the cytotoxicity assay on CHPV infected cells helped to understand the time frame and dynamics of CHPV gene expression (Fig. S1), IFA on these cells allowed us to monitor condensation and punctate localization of viral proteins to form CHPV-IBs (Fig. 1). Although sometimes identified with a different name eg. Negri bodies in Rabies Virus, IBs can be seen consistent among members of *Rhabdoviridae* as well as among negative strand RNA viruses (36, 69, 70). For instance, a closely related member of the family, VSV forms TIA-1+ SG like structures that colocalize with viral replication proteins and RNA (69). Virus infection, in general, inevitably induces metabolic stress and promotes SG production. Thus, we performed this study to understand the course of CHPV-IBs formation and their evasion of SGs; in pursuit of identification of host factors, involved in this regulation. We found that CHPV infection although did not induce formation of distinct SGs but showed association of its IBs with multiple RNA binding proteins of SGs (TIA-1, PAPB1, Ago2, eIF3η, and G3BP)(Fig. 2). We anticipate involvement of more SGPs in CHPV-IBs formation and it would be important to characterize IBs further to get more insight about their composition with respect to cellular proteins. In this direction, whether assembly of IBs is triggered by scaffolding activity of a cellular or a viral component; would be another important question to address in the future investigation. Our previous study on CHPV showing its phosphoprotein mostly intrinsically disordered at its N-terminal might also be a potential factor to be investigated further for this role and in understanding the mechanism of the CHPV-IBs formation.

Our investigation subsequently focused on differentiation of IBs with canonical SGs. Although SGs are often assumed to be uniform entities formed under different stresses, yet their protein and mRNA composition varies (71). The formation of SGs is a dynamic and reversible assembly which disintegrate and cellular translation resume when cell starts to recover from stress. To understand the dynamic properties and to address whether already formed IBs can be strained to disintegrate by limiting the production of its constituents, we first tested CHX to be an inhibitor of CHPV-IBs as already demonstrated earlier for SGs (72). Here, we found CHX can effectively reduce the size and exhibit enforced disassembly of IBs by 3 hrs of its treatment on CHPV infected cells (Fig 3). The inhibitory effect of CHX on IBs not only confirmed IBs to be virally induced but also allowed us to chase the time of their disassembly. When compared to SA-induced conventional SG, IBs were found to be different than SGs in terms of kinetics of their disassembly in the presence of CHX. Altogether, it indicated that IBs are formed through a different process than SG and they might have a different composition and architecture and thus different kinetics of disassembly than conventional SG.

Next, we aimed to understand whether association of TIA-1 is a proviral or antiviral to CHPV replication. TIA-1 is a key player in the formation of SG which in turn triggers the antiviral cellular response and limit virus production (40). However, TIA-1 also exhibits a proviral role in the replication of *Flaviviruses* (73), so its antiviral effect is not universal for all viruses. Next, we sought to determine whether participation of TIA-1 in IBs is proviral or antiviral. Since, PKR is widely accepted to play an antiviral role in infection by many viruses of several families (74) therefore, we postulated that PKR might be a host inhibitory protein to block CHPV production. To our surprise, independent silencing of TIA-1 or PKR in Vero cells showed similar effects of significant decrease in CHPV virion production, and suggested a proviral role of both TIA-1 and PKR (Fig 4). The proviral role of TIA-1 in CHPV replication bear a resemblance to its supportive role in *Flavivirus* genome RNA synthesis through interaction with viral components and inhibiting SGs formation (75). Our observation through IFA on CHPV infected cells also revealed colocalization of PKR with CHPV-IBs, but not with SGs (Fig 5). Here, it may be noted that PKR association with IBs makes them different than SGs in terms of their composition. CHPV resembles to porcine reproductive and respiratory syndrome virus (PRRSV), where PKR has been demonstrated to play a proviral role in viral replication by modulating viral gene transcription (76). It was also demonstrated later that PRRSV selectively inhibits PKR activation to prevent inflammatory response. Interestingly, this inhibition of PKR was shown to be dependent on the viral nsp1β protein to co-optimize G3BP to inhibit PKR activation in viral replication factories (77). It will need further investigation to understand the mechanism of proviral function of both TIA-1 and PKR.

While the process of IBs remains completely unknown, the canonical SG formation is initiated as a consequence of phosphorylation of α subunit in elF2α at a specific serine (Ser 51) residue (78). The phosphorylation of eIF2αacts as a trigger which causes the prion-like aggregation, phase separation and recruitment of TIA-1 to SGs (79).Typically, eIF2α functions as a vital initiation factor in promoting the binding of tRNA^met^ to the 40S ribosome, to promote mRNA translation in a GTP-dependent manner. Different types of stress (oxidative, heat, or nutrient deprivation) can induce eIF2α phosphorylation by activation of four different eIF2α kinases (GCN2, PKR, PERK, and HRI) (56).

To further compare the process of IBs formation with SGs, our investigation focused on the activation of PKR/eIF2α for recruitment of TIA-1 and other RNA binding proteins to CHPV-IBs, as it happens during SGs formation. Here, we found CHPV infection could neither activate PKR nor induce eIF2α phosphorylation and thus suggesting that the recruitment of TIA-1 and other RNA binding proteins to CHPV-IBs occurs independent of activation of this pathway (Fig 6). Our observation on partial inhibition of PKR and eIF2α phosphorylation treated but CHPV infected cells may be speculated about condensed PKR in IBs remaining inactive and inaccessible to PACT-mediated activation during SA treatment. Taken everything together, it can be concluded that CHPV-IBs and SGs are distinct in terms of their disassembly, composition and in the process of their formation. Our proposed model in Fig. 7 shows the two distinct pathways where activation of PKR pathway plays occurs in formation of SGs but not in the CHPV-IBs. PKR in its inactive form however plays an unknown proviral function in the formation of IBs. Moreover, presence of viral proteins (shown in Red) in IBs and presumably viral RNA, make their composition different from SGs. It will however be interesting to know the presence of cellular mRNA in IBs. Altogether, our study not only provides insight in CHPV replication and leaves several important questions to understand CHPV biology but also makes the pavement for designing antiviral therapy against Chandipura Virus.

**Fig. 7.**
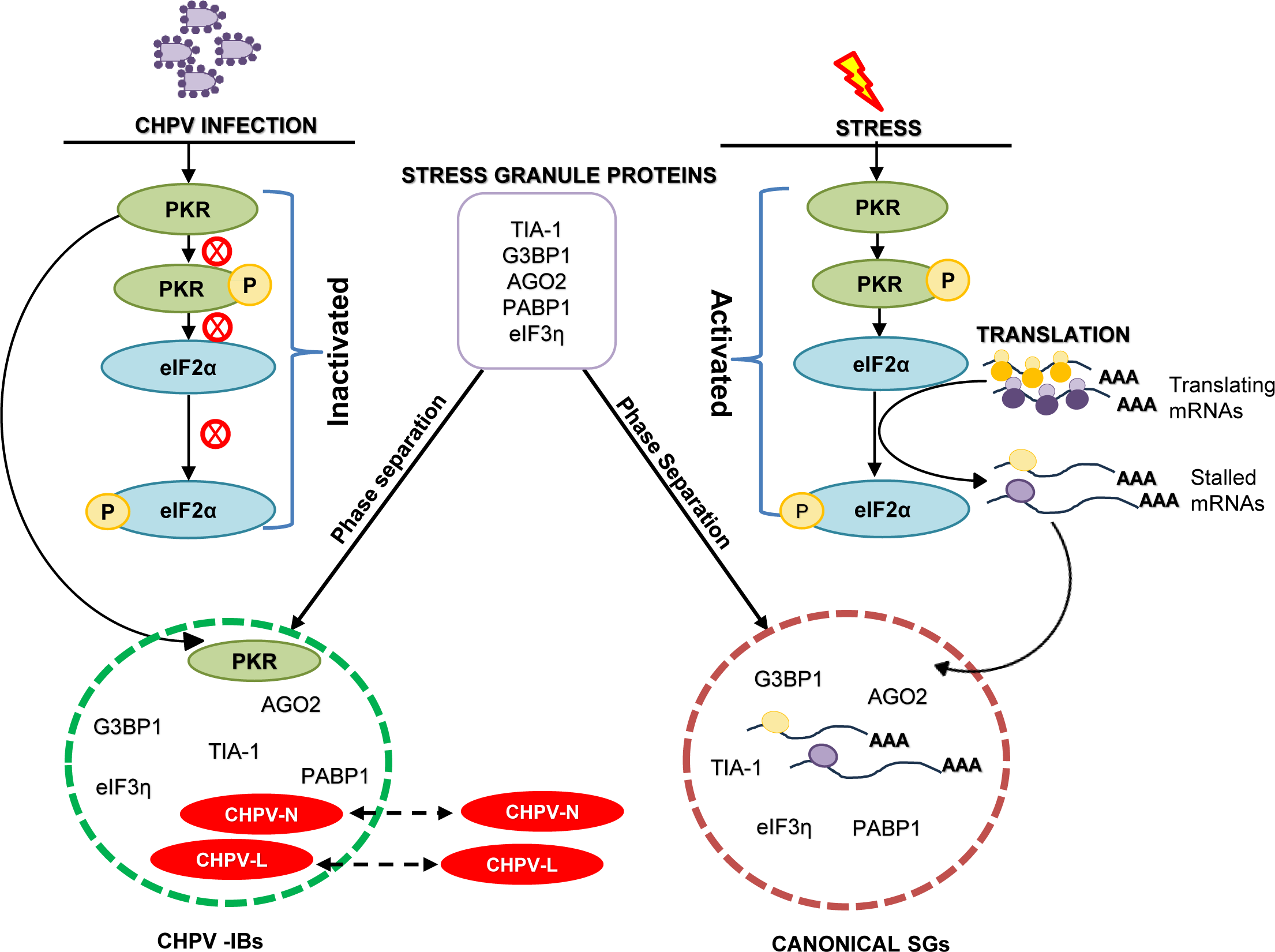
A schematic model showing distinctness in the process of SGs and CHPV-IBs formation. In this model, CHPV proteins (N and L) in the cytoplasm condense and co-localize together in association with other viral, cellular SG proteins, and PKR to form CHPV-IBs. In contrast to formation of SGs, IBs form independent of the activation of PKR/eIF2α phosphorylation.

## Material and Methods

### Material and Methods

#### Cell cultures and virus propagation

Vero E6 Cells (ATCC, Manassas, VA) and N2A cells were cultivated in DMEM (cat no. 12430047, GIBCO/*Thermofisher scientific*) supplemented with 10% heat inactivated fetal bovine serum (cat no. 16140-071, GIBCO/*Thermofisher scientific*). A human patient derived CHPV (strain no. 1653514) was obtained from Dr. Dhrubajyoti Chattopadhyay’s lab (Calcutta University). The virus was propagated in the Vero cells, and viral titer (5×10^7^ pfu/ml), was measured using plaque assay.

#### Antibodies and Chemicals

The antibodies used for this study and their respective working dilutions in Western blotting (WB) or Immunofluorescence assay (IF) are as following. Rabbit polyclonal anti-CHPV-N (1:5000, WB and 1:250 IF) and mouse anti-CHPV-L (IF 1: 250) antibodies were kindly provided by Dr. Dhrubajyoti Chattopadhayay. Mouse anti-TIA-1 (cat no. SC-166247), Rabbit anti-p38 (cat no. SC-728, Goat anti-eIF3η (cat no. SC-16377) were obtained from Santa Cruz Biotechnology. Rabbit anti-elf2α (cat no.9722S), Rabbit anti-Phospho-eIf2α (Ser 51) (cat no. 9721S), Rabbit anti-Phospho-p38 (cat no. 9211S) were obtained from Cell signaling technology. Mouse anti-AGO2 (cat no. 57113, Abcam), mouse monoclonal anti-eIF2AK2 (PKR) (cat no. H000005610-M01, Abnova), Rabbit anti-Phospho-PKR (pThr454) (cat no.527460, Milipore), Mouse anti-PABP1 (cat no. MA1 34079, Invitrogen/*Thermofisher scientific*) and Mouse anti-β-Tubulin (cat no. T5201, Sigma Aldrich) were obtained from their respective companies. The secondary antibodies Alexa flour 568 (H+L) Donkey anti-Goat (cat no. A11057), Alexa fluor 488 (H+L) goat anti-rabbit (cat no. A11008), Alexa fluor 568 (H+L) goat anti-mouse (cat no. A11004), and Alexa fluor 405 (H+L) goat anti-rabbit (cat no. A31556) were obtained from *Thermofisher scientific*.

#### Induction of cellular stress

Sodium arsenite (SA, NaAsO₂, cat no. 1062771000, Sigma Aldrich) solution (0.05M). To induce oxidative stress the cells were cultivated in fresh culture medium containing 1 mM SA for40 min [Sharma et al PLos Path].

#### Immunofluorescence Assay

Adherent Vero E6 cells were grown directly on glass coverslips. Immunofluorescence staining was performed as described previously [Sharma et al Plos Path, NAR, JBC]. Briefly, the pretreated or infected cells were washed with PBS, fixed with 4% paraformaldehyde (PFA), permeabilized with 0.4% Triton X-100 and blocked with 2% BSA (bovine serum albumin, cat no. A9647, Sigma Aldrich) dissolved in Tris-buffered saline containing 0.05% of Tween-20 (TTBS). Primary antibodies diluted in blocking buffer were incubated with slides overnight at 4°C or 3hrs at 37°C in a humidified chamber. Alexa fluor-conjugated secondary antibodies (1: 500, *ThermoFisher Scientific*) were diluted in blocking solution and incubated with slides at 37°C in humidified chamber for 2hrs. The slides were washed atleast 4 times with TTBS and before mounting the cells nuclei were visualized by 5 min counterstaining with wash buffer containing Hoechst dye 33342 (1:10,000 dilution, cat no. B2261, Sigma-Aldrich).

#### Staining of cellular lipid droplets

A stock solution of 10mg/ml (38 mM) BODIPY™ 493/503 (4,4-Difluoro-1,3,5,7,8-Pentamethyl-4-Bora-3a,4a-Diaza-s-Indacene/ cat no. D3922, *ThermoFisher Scientific*) was prepared in DMSO which was diluted in DMSO (1:100) to make as sub-stock of 0.1 mg/ml (0.38 mM). This was further diluted in PBS (2.5 µl in 1 ml PBS) working 1µM to stain neutral lipids in cell.

#### Expression vectors and construction of plasmids

The following vectors were used to express recombinant proteins: A flag-tagged CHPV-N (Chandipura virus-Nucleoprotein) expressing mammalian expression vector was generated by subcloning CHPV-N gene from PET-3a-CHPV-N plasmid (gift from Dr. Dhrubajyoti Chattopadhayay) in pFLAG-CMV6a (Sigma). Primers F-5’ TTTATA AAGCTT ATGAGTTCTCAAGTATTC3’ and R-5’ TTTATA GGATCCTCATGCAAAGAGTTTCCT3’ containing the Hind III and BamHI sites respectively were used to amplify CHPV-N gene. The PCR product was sub-cloned in between sites Hind III and BamHI in MCS of pFLAG-CMV6a plasmid under CMV-promoter. The recombinant plasmid was purified using QIAgen mini prep kit (cat no.27106, Qiagen) as per manufacture’s protocol. The resulting plasmid subsequently named as pFlagCMV6-N (pNRS1), was verified by restriction digestion and sequencing. GFP-G3BP1 wild type construct was kindly provided by Dr. Jomon Joseph (NCCS, Pune, India), originally obtained from Dr Jamal Tazi (Institut de GénétiqueMoléculaire de Montpellier, France). GFP expressing plasmid (pEGFP-N1) was obtained Clontech (cat no. 6085-1).

#### Plasmid transfection

Plasmid transfections were performed using Lipo2000 transfection reagent (cat no. 11688-030, Invitrogen by *ThermoFisher Scientific*) according to the manufacturer’s instruction. Unless indicated, for IFA and Western analysis, Vero E6cells (3 ×10^5^) were plated a day prior to which 1 µg of plasmid DNA transfection in a 6-well plate was performed.

#### siRNA transfection and measurement of CHPV virion production

For siRNA mediated silencing, ∼1.5×10^5^Vero E6 cells growing in a 12-well plate were transfected twice, respectively, at an interval of 24 hr with siRNAs targeting human PKR/EIF2AK2 (Assay ID 142330-*Thermofisher scientific*) or human TIA-1 (Assay ID 139893-*Thermofisher scientific*) or GFP targeting siRNA as a negative control (cat no. P-002048-01-20, Dharmacon) using Lipofectamine 3000 transfection reagent (cat no. L3000-008-Invitrogen*/ Thermofisher scientific*). Total cell extract was collected 24hrs after the second siRNA transfection to measure the knockdown efficiency by western blotting.

Alternatively, for CHPV virion production and titration assays, Vero cells in a 12-well plate were transfected with siRNAs targeting human PKR or TIA-1 and after 20 hr of second round of siRNA transfections as described above, were infected with 0.01 MOI CHPV in 1 ml of DMEM medium. Culture supernatants were harvested 18hr after infection, cleared by centrifugation at 2000 rpm for 10 min, and aliquoted for further measurement of viral titer by plaque assay. In parallel, three plates of siRNA transfected cells, 1 MOI CHPV was incubated in 1 ml of complete medium. In one of the plates, Vero cell extract were prepared after 6hr of infection as mentioned below for western blotting to detect viral CHPV-N or cellular PKR or TIA-1, p-eIF2α, total eIF2α, and tubulin using specific antibodies. In another plate, after 6hr of infection, Vero cells were harvested for RNA extraction to measure amount of viral RNA. In yet another parallel plate of siRNA transfected cells, 1 MOI CHPV was incubated in 1 ml of complete medium. After 4hrs infection, Vero cells were fixed using 4% PFA and later immunostained for CHPV-N or cellular PKR or TIA-1.

#### Plaque Assay

Vero cells (∼1.5×10^5^ cells/ well) were grown in 12 well plate and allowed to grow for 48 hr to reach confluency of ∼90-95%. Cells were washed twice with 1X PBS and added with cell free supernatant containing CHPV virions diluted in serum free medium and incubated for 2hr at 37°C in CO_2_ incubator. Again, cells were washed with PBS twice and overlayed with 2x DMEM mixed with equal volume of 2% low melting agarose. Plates were then incubated for 18-24 hrs at 37°C in CO_2_ incubator. To visualize plaques, cells were stained with crystal violet (0.5% W/V crystal violet in 25% methanol) for 2 hrs and the agarose overlay was discarded. Finally, wells were rinsed with water for visual counting of the plaques.

#### Western blotting

Unless indicated otherwise, protein samples for Western blot were prepared by direct lysis of the cells in 2X SDS sample buffer (0.02% bromophenol blue(w/v) in 4%SDS, 120mM Tris HCl pH=6.8 and 20% glycerol(v/v)) containing 5% 2-mercaptoethanol (Sigma-Aldrich). Samples were resolved on a 10% SDS-PAGE gel in 1 × Tris glycine SDS buffer. The signal was detected with Chemiluminescent Substrate (BIORAD,cat no.1705061).

#### RNA extraction, cDNA preparation and qRT-PCR

Vero-E6 cells growing in 12 well plate after siRNA transfection and/or infection with CHPV were lysed in 1ml Trizol (cat no.15596026, Invitrogen/ *Thermofisher scientific*), 200μl of chloroform was added and mixed vigorously for 15 sec and incubated at RT for 2-3 mins. The mixture was centrifuged at 13000rpm for 15 min at 4°C to separate out aqueous and phenol layer. The aqueous layer was separated carefully; equal amount of isopropanol was added and incubated on ice for 30 mins to precipitate the RNA. Precipitated sample is centrifuged at 13000rpm for 15 min at 4°C. The RNA pellet was washed using 70% ethanol and then resuspended in 15μl of nuclease free water. DNase treatment was given to the extracted RNA for 15 mins at RT using DNase-1 kit (cat no.18068-015, Gibco/*Thermofisher scientific*) according to manufacturer’s protocol. The DNA free RNA was used for cDNA preparation using Takara first strand cDNA synthesis kit (cat no.6110A, Takara Bio Inc., Japan) according to manufacturer’s protocol. The RNA template of ∼1µg was incubated with oligo dT primers (50µM) and dNTP mixture(10µM) at 65°C for 5mins and then 5mins on ice. Later the template RNA/ Primer mixture was combined with 5X Prime script buffer, 20U of RNase inhibitor and 200U of Prime script reverse transcriptase were set to PCR (30°C for 10mins, 42°C for 60min and 72°C for 15mins) to synthesize first strand complementary DNA. The prepared cDNA (0.25 µl) were used for quantitative real time-PCR using 2x sybr green master mix (cat no.4344463, applied biosystems/*Thermofisher scientific*) to measure the amount of CHPV-N specific mRNA in infected Vero cells using CHPV-N specific primers (forward 5′-ACCTGGCTCCAAATCCAATAC-3′ and reverse 5′-GGTGGATCAGACGGAGAGATA-3′). β-actin (forward 5′-GACAGGATGCAGAAGGAGAT-3′ and reverse 5′-GCTTGCTGATCCACATCTGC-3′) was used as the housekeeping gene.

#### Cycloheximide treatment

To understand effect of CHX on SGs formation, Vero cells were grown on coverslips in three 35mm dishes parallelly. One of the dish was pre-treated with 100 µg/ml of CHX for 1hr; later media was replaced with complete media containing 1mM SA for 40min in presence of CHX. Another dish was treated only with 1mM SA for 40min. Yet another dish was not treated at all and served as a control. All the 3 plates were fixed with 4%PFA and stained for TIA-1 and PABP1 using their respective antibodies.

To understand the effect of CHX on CHPV-IBs, Vero E6 cells were grown on coverslips in 6 well plates in two sets and allowed to reach confluency of 70-80%. Cells were infected with 2 MOI CHPV and after 3 hrs of infection; virus containing media was replaced with complete media containing 100 µg/ml of CHX. After 1hr of CHX treatment, cells were fixed with 4% PFA and used for IFA. Another set of cells were infected with 2 MOI CHPV and after 1 hr of infection, virus containing media was replaced with complete media containing 100 µg/ml of CHX. After 3 hr of CHX treatment cells were fixed with 4% PFA and used for immunofluorescence staining.

To understand the dynamics of CHPV-IBs in contrast to SGs through inhibition of protein synthesis by cycloheximide (CHX), Vero E6 cells were grown on coverslips in 35mm dishes and allowed to reach confluency of 70-80%. Cells were infected with 2 MOI CHPV, after 4hrs of virus infection cells were treated with 100 µg/ml of CHX for 15mins, 30mins, 1hr and 2hrs respectively. Then fixed with 4% PFA and used for immunofluorescence staining. Another set of cells were first exposed to 1mM of SA for 40 min to form canonical stress granules and then treated with 100 µg/ml of CHX for 15 mins, 30 mins, 1 hr and 2 hrs respectively. Then cells were fixed with 4% PFA and used for immunofluorescence staining.

#### MTT assay

To understand the metabolic state of cells upon CHPV infection MTT assay was used. Vero cells were plated in 96 well plate and allowed to reach confluency of 70-80%. Cells were infected with 5 MOI CHPV at different time points; after completion of virus infection virus containing media was replaced with media containing 5 µg/ml MTT reagent (Thiazolyl blue tetrazolium bromide, cat no.298-93-1, Gold biotechnology). Incubate at 37°C for 3-4 hr and then remove MTT containing media and add 100 µl of DMSO to each well to dissolve the formazan crystals formed, after 15 mins measure OD at 594nm. The amount of formazan formed is directly co-related to live cells in the well.

#### Confocal imaging and Statistical analysis

Fluorescence images were captured with Leica SP8 confocal microscope equipped with 63X oil immersion objective lens. The images were processed on Las X free software.

Data were represented as mean of 3biological replicates ± standard deviation from the mean. Student’s T-test was performed on Graph pad Prism8 to evaluate the statistical significance. A p-value of <0.05 was considered statistically significant.

Quantification of CHPV-IBs were performed on image J. To count and quantify CHPV-IBs the “Analyze Particle” plugin of image J was used for which images were converted to 8 bit gray-scale, scale was set to 10μm and threshold was set to eliminate background and select only CHPV-IBs. Each cell was then marked using “free hand tool” and analyze particle plugin was applied to analyze number of CHPV-IBs per cell and size of each CHPV-IB inside the cell. To calculate the Pearson’s co-relation analysis of co-localizing IBs, TIFF images of red and green channel were opened in ImageJ individually and merged. The merged image was converted to RGB color format, colour threshold was set to select only co-localizing IBs. Total co-localizing area was measured and JACop plugin of ImageJ to get Pearson’s co-relation coefficient.

## Acknowledgments

We thank Prof. Dhrubajyoti Chattopadhyay for providing us the plasmid, antibodies and Chandipura virus (strain 1653514). We also thank Dr. Jomon Joseph for a plasmid expressing GFP-G3BP1. SS is a recipient of junior research fellowship from Department of Biotechnology (DBT, DBT/2019/IGIB/1203). NRS received financial support from the Ramalingaswami Re-entry fellowship from the Department of Biotechnology (DBT), Ministry of Science and Technology, Government of India (BT/RLF/Re-entry/40/2018).

## Supplementary figures

**Fig. S1A.**
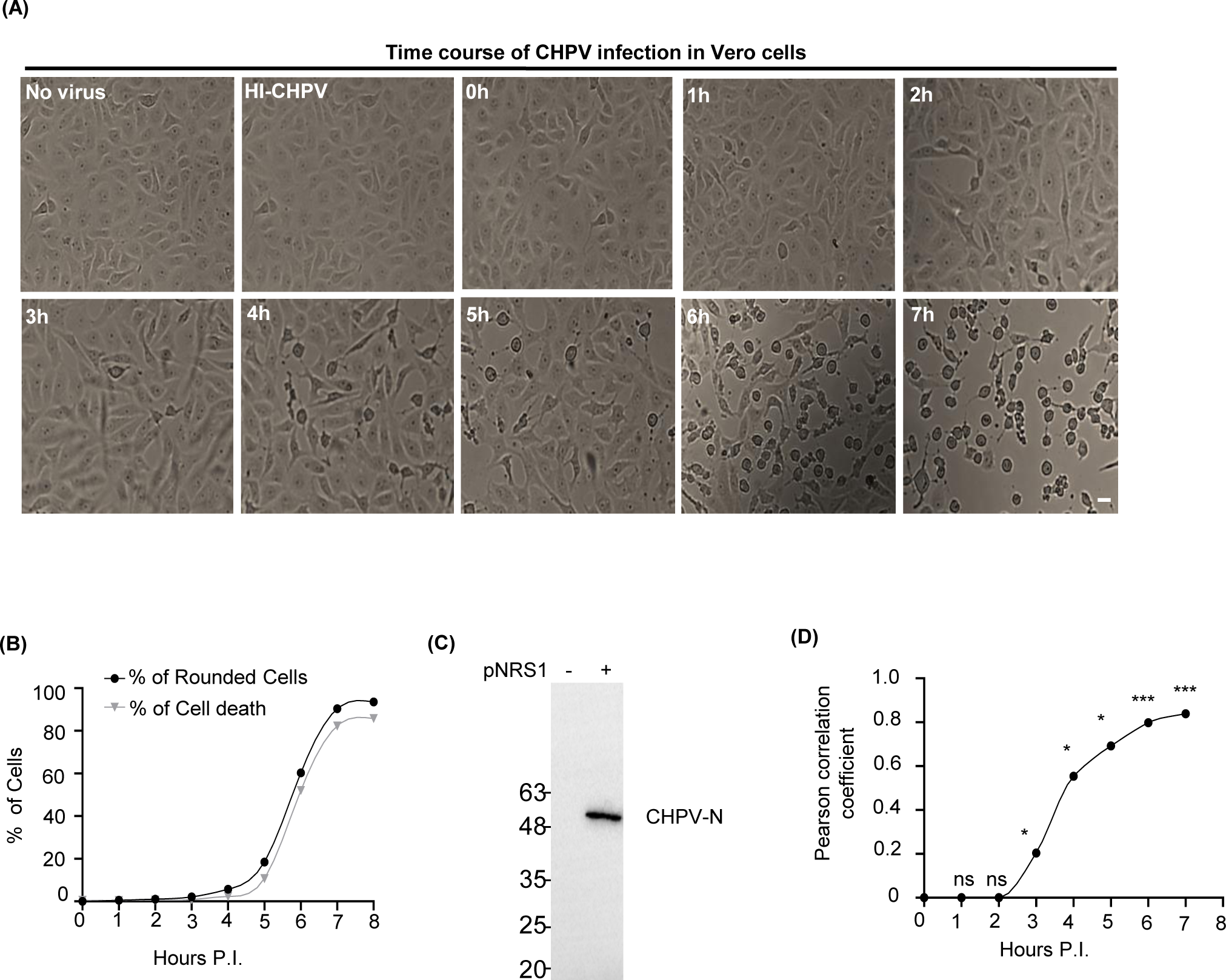
Time course of CHPV infection. Vero Cells were infected with 5MOI live CHPV for indicated time or with heat-inactivated (56°C/20 min) CHPV (HI-CHPV) for 7 hrs. Cells were observed under a bight field microscope for cytopathic changes and images were collected. Same cells were further immunostained with anti-CHPV-N and anti-CHPV-L antibodies. Scale bar = 20 µm. **Fig. S1B** Graph showing percentage of cells in Fig. S1A with round morphology and percentage of cell death evaluated by MTT assay, with mean ± SD from three independent experiments. **P<0.01 in Chi-squared test. **Fig. S1C** Western blot detection of the ectopic expression of flag-tagged CHPV-N protein using a rabbit polyclonal anti-Flag antibody. **Fig. S1D** Graph showing co-relation between CHPV-N and CHPV-L puncta in immunostaining (Fig. 1B) by plotting Pearson’s co-relation coefficient for each time point. (n=3)

**Fig. S2A.**
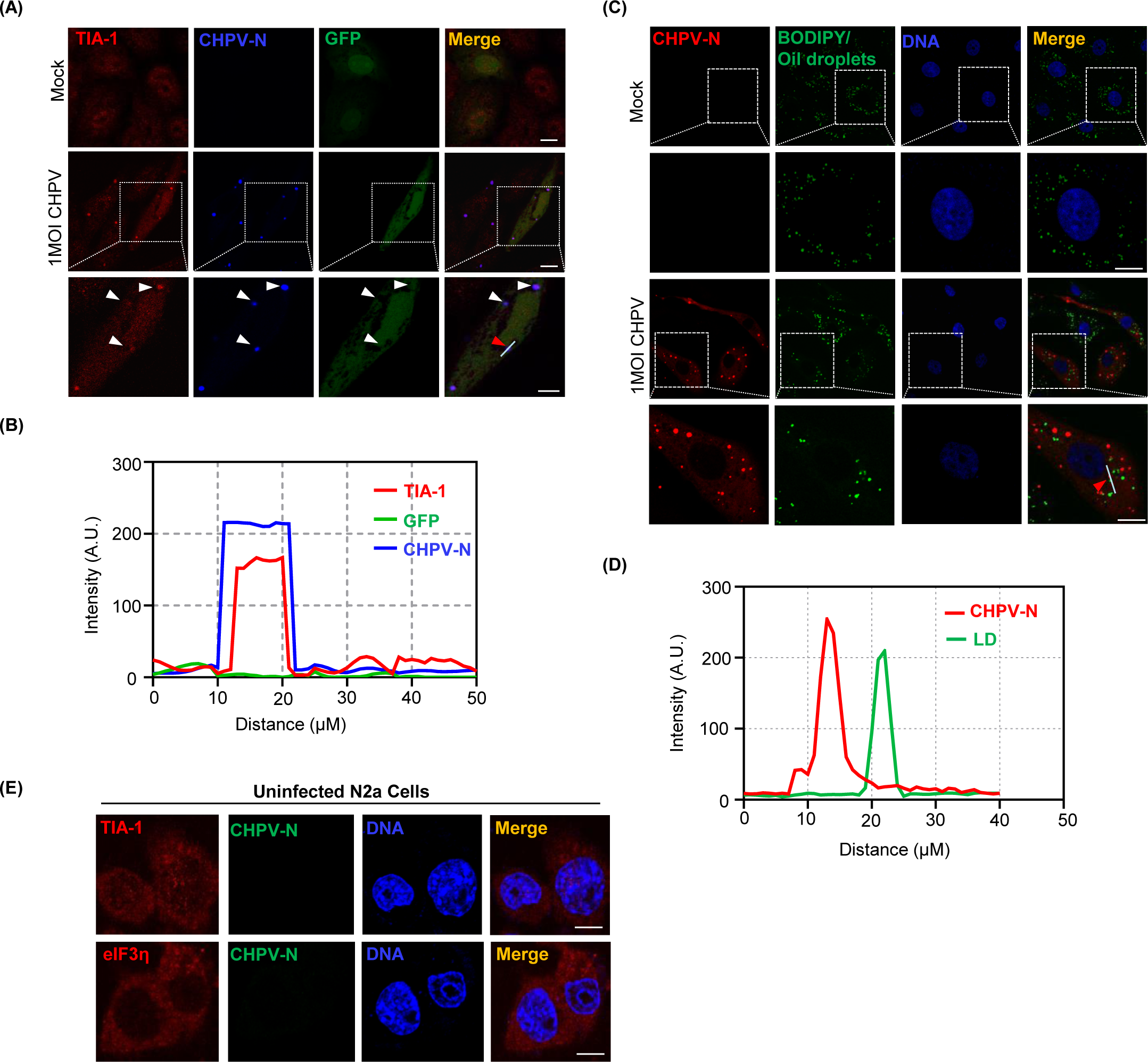
Absence of co-localization between CHPV-IBs and ectopically expressing GFP mock plasmid. Vero cells were transfected with GFP alone vector and after 24 hrs, infected with 1 MOI CHPV for 4 hrs. Cells were then fixed with 4% PFA and immunostained for CHPV-N (blue) and TIA-1 (Red), showing GFP does not co-localize with the IBs in contrast to GFP-G3BP1 which co-localizes with CHPV-IBs (Fig 2J). Scale bar = 10 µm. **Fig. S2B** Graph shows the distribution of signal intensities in arbitrary units (A.U.) for TIA-1, GFP and CHPV-N across a line indicated by red arrow, using the Image J software. **Fig. S2C** Absence of Co-localization between lipid droplets (LDs) and CHPV-IBs. Vero cells were infected with 1 MOI CHPV for 4 hrs. Cells were then fixed with 4% PFA and immunostained for LDs using BODIPY D493 stain and CHPV-N antibody. **Fig. S2D** Graph shows the distribution of signal intensities in arbitrary units (A.U.) for LDs and CHPV-N across a line indicated by red arrow, using the Image J software. **Fig. S2E** Uninfected N2A cells stained with TIA-1 (red, upper panel) and eIF3η (red, lower panel) to serve as background control for Fig. 2L. The nuclei were counterstained with Hoechst dye. Scale bar = 10 µm

**Fig. S3A.**
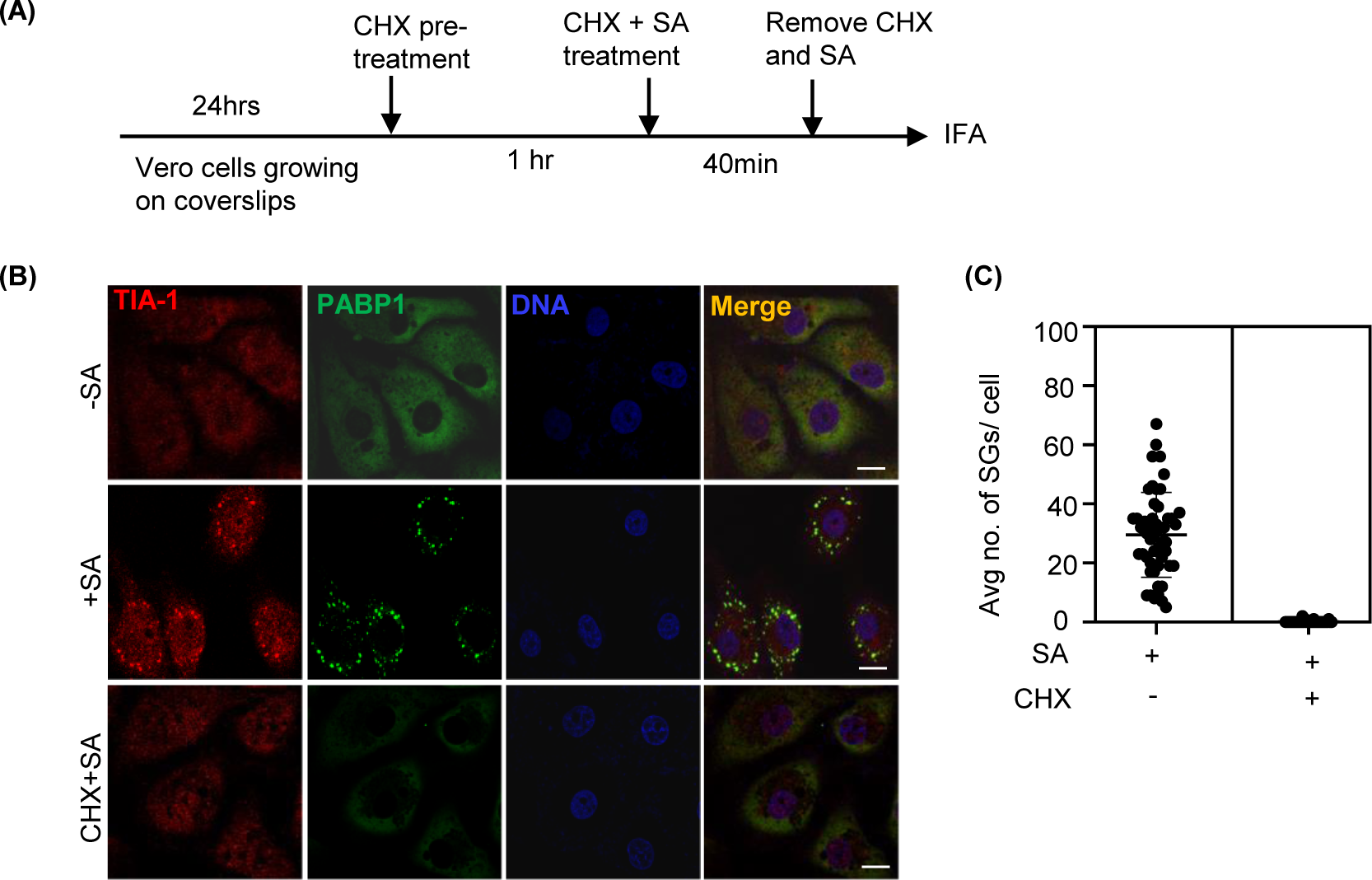
The schematic shows outline of experimental design. Vero cells grown on coverslips pretreated with Cycloheximide (CHX) 50 μg/ml for 1hr. Both untreated and treated cells were incubated with 1mM SA 40 min in the absence or presence of CHX respectively. **Fig. S3B** Cycloheximide (CHX) treatment inhibits SA-induced SGs. Vero cells were left untreated or pretreated with CHX (50 μg/ml) for 1 hr. Both untreated and treated cells were incubated with 1mM SA for 40 min in the absence or presence of CHX respectively were fixed with 4% PFA and immunostained for TIA-1(red) and PABP1 (green). The nuclei were counterstained with Hoechst dye. Scale bar = 10 µm. **Fig. S3C** Graph (Fig. S3C) shows average number of SGs formed per cell by SA treatment (∼50 SGs/cell) versus almost no SGs formed upon CHX pre-treatment. (n=50)

**Fig. S4A.**
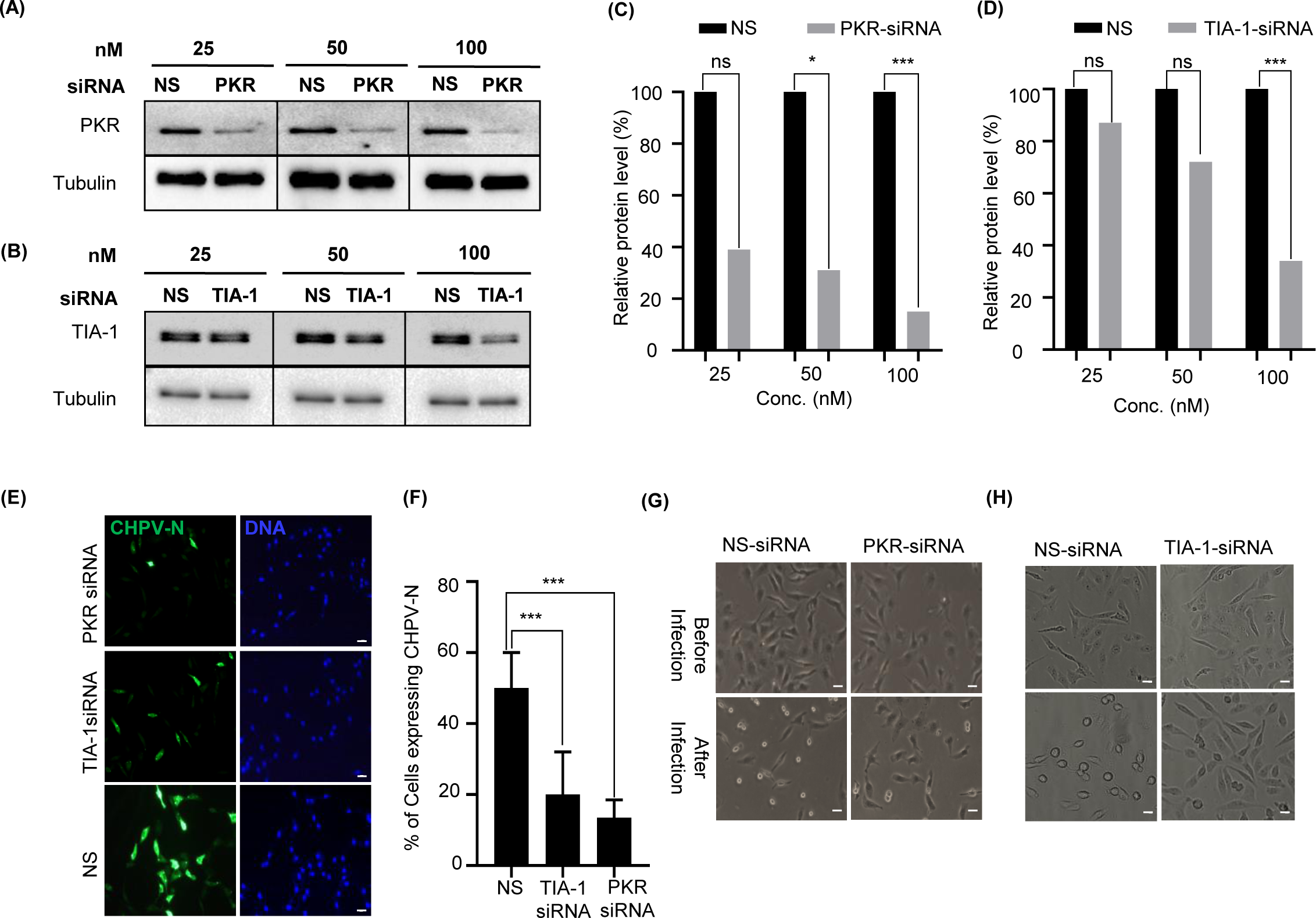
Dose optimization of siRNA knockdown of PKR. Vero cells were double transfected with increasing doses of PKR siRNA (25 nM, 50 nM and 100 nM) to optimize efficient knockdown of ∼70-80%. **Fig. S4B** Dose optimization of siRNA knockdown of TIA-1. Vero cells were double transfected with increasing doses of TIA-1 siRNA (25 nM, 50 nM and 100 nM) to optimize efficient knockdown of ∼70-80%. **Fig. S4C-D** Graphs in showing relative intensity of PKR and TIA-1 after knockdown by respective siRNAs and by normalizing with tubulin as loading control. **Fig. S4E** Cellular CHPV-N expression is reduced upon siRNA mediated knockdown of TIA-1/PKR. Vero cells were double transfected with TIA-1/PKR siRNA, infected with 1MOI CHPV for 4 hrs and fixed with 4% PFA for immunostaining. The cells were immunostained for CHPV-N and nuclei were counterstained with Hoechst dye. Scale bar = 20 µm. **Fig. S4F** Quantification of CHPV-N expression upon siRNA mediated knockdown of TIA-1/PKR. The graph shows significant decrease in percentage of cells expressing CHPV-N after knockdown of TIA-1/PKR (n=3). **Fig. S4G-H.** The phase contrast images showing the inhibition of CHPV-induced cytopathic effect (round cell morphology) in Vero cells after siRNA knockdown of TIA-1 or PKR (lower panel, after infection). Before CHPV infection, cells were completely healthy after double transfection of both siRNAs (upper panel, before infection). Scale bar = 20 µm.

**Fig. S5A.**
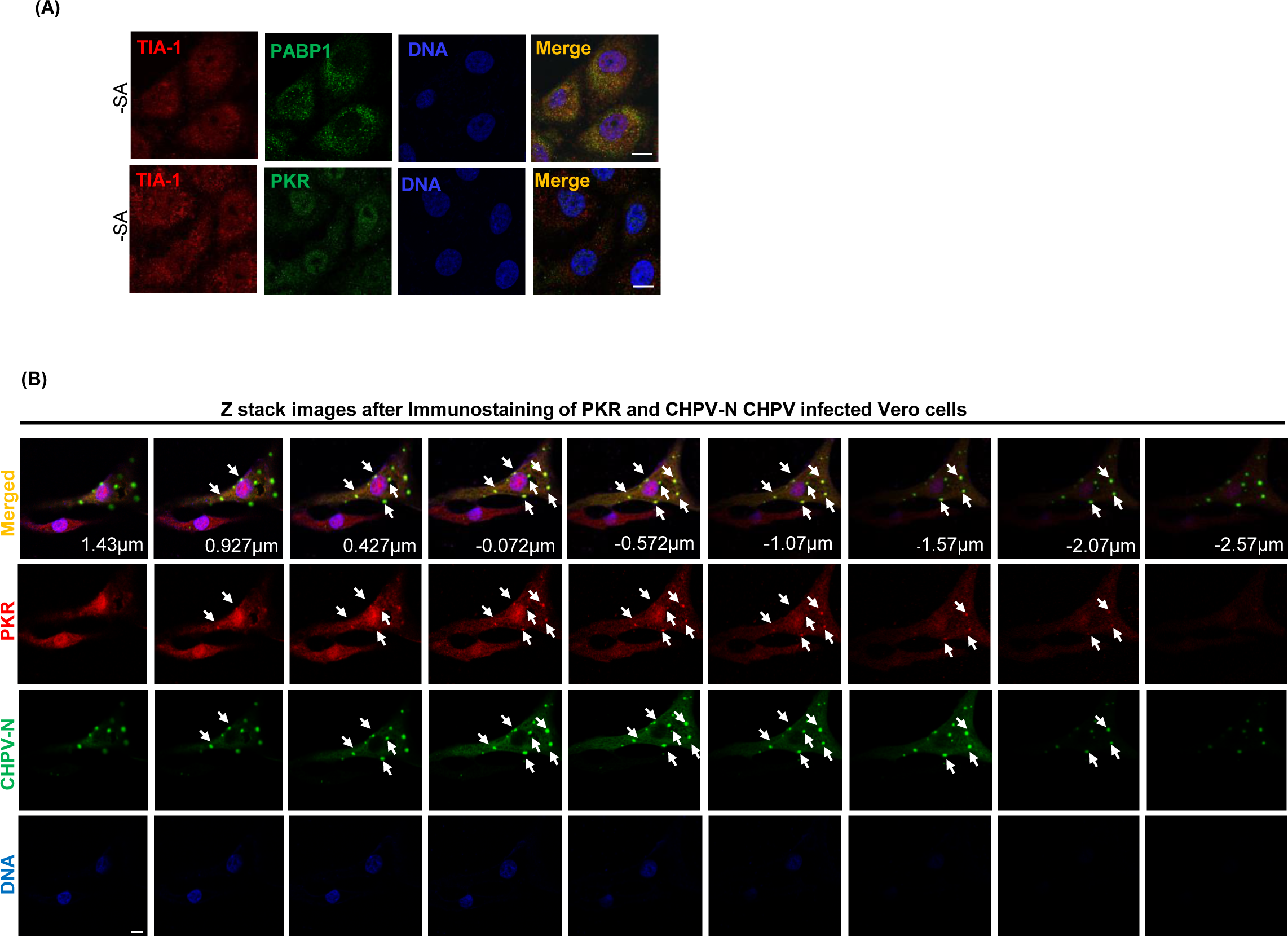
Vero cells were immunostained without SA treatment to observe the background staining for TIA-1 (red) and PABP1 (green) to serve as control for Fig 5A. **Fig. S5B** Vero Cells were infected by CHPV 1MOI for 4 hrs to induce IBs. Cells were fixed with 4% PFA and then processed for IFA to detect CHPV-N/ PKR for CHPV-IBs. The nuclei were counterstained with Hoechst dye. Scale bar = 10 µm. Confocal z-stack was used to generate the maximum intensity projection of all three channels with 9 sections (from 1.43µm to −2.57µm) of images, showing the PKR localizes with CHPV-N puncta.

**Fig. S6A.**
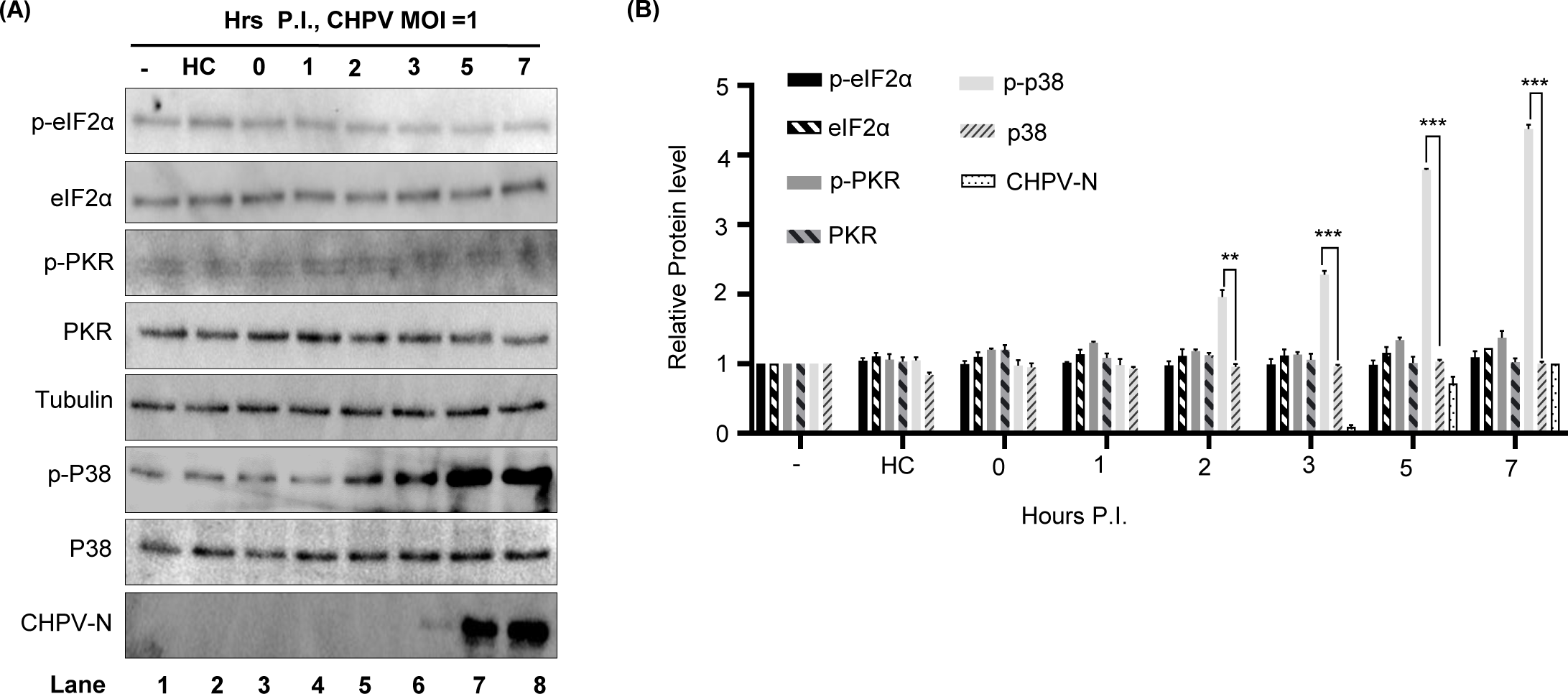
Kinetics of p38 activation with no change in p-PKR/p-eIF2α. Vero cells were infected with live CHPV (1 MOI) for the indicated time. Infection with heat-inactivated (HI) CHPV was done for 7 hrs. The phosphorylated forms of eIF2α (p-eIF2α, Ser51), PKR (p-PKR) and p38 (p-p38) were measured by Western blot analysis using phospho-specific antibodies. Total level of these kinases was determined by a pan-antibody. Tubulin served as a loading control. **Fig. S6B:** Graphical representation of the relative amount of p-eIF2α, eIF2α, p-PKR, PKR, p-p38 and p38 protein in each sample after normalizing to tubulin was plotted over the no virus control in lane 1 set as 1. The error bar indicates mean ±SD (n=3).

**Fig. S1 video:** A 3-D video rotation of confocal z-stack of Vero cells infected with CHPV 1MOI for 4 hrs was reconstructed. The cells were immunostained with CHPV-N and PKR specific antibodies. The nucleus was counterstained with Hoechst dye. Confocal z-stack was used to generate the maximum intensity projection of all three channels, showing PKR co-localizes with CHPV-N in x, y and z axis.

**Fig. S2 video:** Vero cells infected with CHPV 1MOI for 4 hrs. The cells were immunostained with CHPV-N and PKR specific antibodies. The nucleus was counterstained with Hoechst dye. A video from 9 confocal z-stack images were reconstructed using las X software along the Z-axis, shows PKR co-localizes with CHPV-N puncta.

